# Manipulations of central amygdala neurotensin neurons alter the consumption of ethanol and sweet fluids in mice

**DOI:** 10.1101/245274

**Authors:** María Luisa Torruella-Suárez, Jessica R. Vandenberg, Elizabeth S. Cogan, Gregory J. Tipton, Adonay Teklezghi, Kedar Dange, Gunjan K. Patel, Jenna A. McHenry, J. Andrew Hardaway, Pranish A. Kantak, Nicole A. Crowley, Jeffrey F. DiBerto, Sara P. Faccidomo, Clyde W. Hodge, Garret D. Stuber, Zoé A. McElligott

## Abstract

The central nucleus of the amygdala plays a significant role in alcohol use and other affective disorders; however, the genetically-defined neuronal subtypes and their projections that govern these behaviors are not well known. Here we show that neurotensin neurons in the central nucleus of the amygdala of male mice are activated by *in vivo* ethanol consumption and that genetic ablation of these neurons decreases ethanol consumption and preference in non-ethanol dependent animals. This ablation did not impact preference for sucrose, saccharin, or quinine. We found that the most robust projection of the central amygdala neurotensin neurons was to the parabrachial nucleus, a brain region known to be important in feeding behaviors, conditioned taste aversion, and alarm. Optogenetic stimulation of projections from these neurons to the parabrachial nucleus is reinforcing, and increases ethanol drinking as well as consumption of sucrose and saccharin solutions. These data suggest that this central amygdala to parabrachial nucleus projection influences the expression of reward-related phenotypes and is a novel circuit promoting consumption of ethanol and palatable fluids.

## Introduction

The central nucleus of the amygdala (CeA) is a heterogeneous structure that plays an important role in the regulation of appetitive, aversive, and ethanol-mediated behaviors (Mahler and Berridge, 2009; Tye et al., 2011; Robinson et al., 2014; McCall et al., 2015; Warlow et al., 2017; Kim et al., 2017; Douglass et al., 2017; Hardaway et al., 2019; Salling et al., 2016). While some data have shed light on neuronal subpopulations influencing fear- and feeding-related behaviors in the CeA (Haubensak et al., 2010; Cai et al., 2014; Douglass et al., 2017), it remains unclear which CeA subpopulations and efferents influence ethanol consumption, particularly during early ethanol seeking (Gilpin et al., 2015; de Guglielmo et al., 2019). A promising CeA subpopulation that may regulate ethanol behaviors are the neurons that express the 13 amino-acid neuropeptide neurotensin (NTS).

NTS is expressed throughout the mammalian brain, including but not limited to the lateral hypothalamus (LH), amygdala, hippocampus, and rostral medulla (Schroeder et al., 2019). Considerable evidence suggests that NTS signaling is critical for reward and anxiety processes (Cáceda et al., 2006; Leinninger et al., 2011; Fitzpatrick et al., 2012; Prus et al., 2014, 2014; McHenry et al., 2017), and global manipulations of NTS signaling disrupt ethanol-related phenotypes (Lee et al., 2010, 2011). However, the roles of individual NTS-positive (NTS+) neuronal populations are not well understood, as the majority of studies investigating NTS+ cells have focused on the LH to ventral tegmental area (VTA) pathway, and particularly on NTS/dopamine interactions (Binder et al., 2001; Leinninger et al., 2011; Kempadoo et al., 2013; McHenry et al., 2017). NTS+ neurons in the CeA (*NTS*^CeA^) have yet to be extensively studied and are in a compelling anatomical and functional position to influence ethanol consumption. Furthermore, early studies identified *NTS*^CeA^ cells that project to the parabrachial nucleus (PBN; Moga and Gray, 1985), a brain region important for fluid consumption.

The PBN, a heterogeneous nucleus that has long been recognized as a sensory relay for taste information, plays a crucial role in the development of conditioned taste aversion (Grigson et al., 1998; Carter et al., 2015). Interestingly, intraperitoneal injections of ethanol induce Fos activation in the PBN (Chang et al., 1995; Thiele et al., 1996). This suggests that the PBN may either be a direct locus for the pharmacological effects of ethanol, and/or receive information regarding the interoception of ethanol. The PBN is also linked to general fluid intake (Edwards and Johnson, 1991) and recent work has identified the PBN oxytocin receptor (*Oxtr1*)-containing neurons as an important locus for fluid satiation (Ryan et al., 2017). An additional subpopulation of PBN neurons, the calcitonin gene-related peptide (CGRP) neurons, are part of an important circuit implicated in suppressing both food and fluid intake (Carter et al., 2013; Ryan et al., 2017). An *Htr2a* CeA-to-PBN (serotonin receptor 2a, *Htr2a*^CeA➔PBN^) projection promotes feeding, suggesting the possibility of a CeA-to-PBN projection that promotes drinking (Douglass et al., 2017). A number of systems have been suggested as a link between food and ethanol consumption such as neuropeptide-Y (NPY; Kelley et al., 2001; Gilpin et al., 2004) and ghrelin (Leggio, 2010). Fluid-consumption related circuits, however, have yet to be examined in this fashion.

To investigate the complex relationship between the CeA and PBN, and better understand the role of the *NTS*^CeA^ neuronal subpopulation in ethanol consumption and appetitive behaviors, we utilized NTS-IRES-Cre mice (Leinninger et al., 2011) in conjunction with region-directed genetic lesion, *Fos* activation, terminal field optogenetic stimulation, and behavioral assays. We find that *NTS*^CeA^ neurons are activated by, and promote ethanol consumption. Furthermore, stimulation of the *NTS*^CeA➔PBN^ projection is reinforcing, and increases the consumption of palatable fluids such as ethanol, sucrose, and saccharin solutions, without altering consumption of neutral or aversive fluids. These data implicate the *NTS*^CeA➔PBN^ circuit as a critical node for the consumption of rewarding and/or palatable fluids.

## Materials and Methods

### Subjects, stereotaxic surgery, virus injection and fiber implantation

#### Mice

All procedures were conducted in accordance with the Guide for the Care and Use of Laboratory Animals, as adopted by the NIH, and with approval of an Institutional Animal Care and Use Committee at UNC-Chapel Hill. Adult male mice 10 weeks and older (>22g) were used for all experiments. C57BL/6J mice were used for the *in situ* tastant exposure experiment (Jackson Laboratories, Bar Harbor, ME). We used adult male NTS-IRES-Cre mice (Leinninger et al., 2011) partially backcrossed onto a C57BL/6J background for all other experiments (Jackson Laboratories, Bar Harbor, ME). Animals were maintained on a reverse 12 hour light cycle with lights off at 7 AM and had *ad libitum* access to food and water (unless noted).

#### Surgery

Mice were anesthetized with inhaled isoflurane (1-3%) and placed in a stereotaxic frame (David Kopf, Germany). For all experiments coordinates for the CeA were as follows (from Bregma, in mm: ML: ± 2.95, AP: - 1.1, DV: - 4.8, for the PBN: ML ± 1.4, AP: -5.4, DV: -4.0 (optical fibers). 300 nL of AAV5-Ef1α-FLEX-taCasp3-TEVp (denoted as: CeA*^NTS^*::casp), AAV5-Ef1α-ChR2-eYFP (denoted as: *NTS*::ChR2 or *NTS*^CeA➔PBN^::ChR2), AAV8-eF1a-DIO-iC++-eYFP (denoted as: *NTS*::IC++ or *NTS*^CeA➔PBN^::IC++), or AAV5-Ef1α-eYFP (denoted as: *NTS*::eYFP or *NTS*^CeA➔PBN^::eYFP) was infused into the CeA at a rate of 100 nL/min. Optical fibers were constructed as previously described (Sparta et al., 2011). Mice were allowed to recover for at least 4 weeks prior to experimentation (8 weeks for optogenetic experiments) to ensure adequate expression of virally encoded genes, and lesioning of target neurons, or protein incorporation into the membrane. All viruses were made by the UNC Viral Vector Core (Chapel Hill, NC) or the Stanford Viral Vector (Palo Alto, CA). Following behavioral studies, animals with ChR2-eYFP construct were perfused, and brains were sliced to verify expression of virus. Animals with no viral expression in either CeA were removed (n=1), while animals with either bilateral or unilateral viral expression were included in the analysis as our pilot data indicated that unilateral expression of the virus was sufficient to drive real-time place preference (RTPP) behavior (data not shown). Animals expressing the caspase construct were euthanized, and brains were flash frozen for validation using fluorescent in situ hybridization (FISH, see below) and compared to their eYFP controls.

### Fluorescent *in situ* hybridization

#### CeA transcript expression

Mice were anesthetized (isoflurane), decapitated, and brains were flash frozen on dry ice. 12 µm slices were made using a Leica cryostat (CM 3050S, Germany). FISH was performed using probes constructed against *Crh*, *Crhr*1, *Pdyn* (type-6, fast blue) and *Nts* (type 1, fast red) and reagents in the View RNA kit (Affymetrix, Santa Clara, CA). FISH was also performed for *Fos* (Mm-Fos-C1, Mm-Fos-C2), *Sst* (Mm-Sst-C2), *Pkcδ* (Mm-Prkcd-C2), and *Nts* (Mm-Nts-C1, Mm-Nts-C2) using the RNAscope Fluorescent Multiplex Assay (Advanced Cell Diagnostics, Hayward, CA). Slides were counterstained with DAPI.

#### In vivo tastant exposure

Singly-housed C57BL/6J mice were habituated to the animal facility for at least 2 weeks. Each animal had homecage access to a single bottle of either water, 6% (w/v) ethanol, 1% (w/v) sucrose, 0.003% (w/v) saccharin or 100μM quinine for 2 hours for 4 consecutive days. On the 5^th^ day, animals had 1 hour of exposure to the same bottle. Half an hour after the bottle was removed, the animals were euthanized for *Nts/Fos* double FISH using RNAscope Fluorescent Multiplex Assay (Advanced Cell Diagnostics, Hayward, CA). CeA slices were taken from approximately bregma -0.8 to -1.9 mm. Experimenters were blinded to consumption conditions for *Fos* and *Nts* counting.

### Immunohistochemistry

As previously described (Pleil et al., 2015), mice were perfused with 4% paraformaldehyde (in 0.01 M PBS), brains were removed and remained in fixative for 24 hours followed by cryoprotection in 30% sucrose/PBS. Subsequently brains were sliced at 40 µm using either a CM 3050S or a VT1000 (Leica, Germany). Sections were incubated overnight at 4°C in blocking solution containing primary antibody – sheep anti-tyrosine hydroxylase 1:500 (Pel Freeze), rabbit anti-neurotensin 1:500 (ab43833, Abcam). The following day, sections were incubated in fluorescence-conjugated donkey anti-rabbit IgG Alexa Fluor 647 secondary antibody (1:800, Jackson Immuno) and donkey anti-sheep 488 (1:200, Invitrogen) for 2 hr in darkness. 435 neurotrace or DAPI was used as a counterstain.

### Microscopy

Images were collected and processed on a Zeiss 710, 780 or 800 a using 20X/0.8 objective and the Zen software (Carl Zeiss, Germany). Image J/Fiji was used for cell counting and data analysis.

### Slice preparation and whole-cell electrophysiology

As previously described (Pleil et al., 2015), animals were anesthetized (isoflurane or pentobarbital/phenytoin) and decapitated. Brains were removed and sliced at a thickness of 200 µm (CeA or PBN) or 300 µm (CeA) using a Leica VT1200 or VT1000 (Germany) in ice-cold high-sucrose low Na+ artificial cerebral spinal fluid (aCSF in mM: 194 sucrose, 20 NaCl, 4.4 KCl, 2 CaCl_2_, 1 MgCl_2_, 1.2 NaH_2_PO_4_, 10 glucose, 26 NaHCO_3_) that had been oxygenated (95% O2, 5% CO2) for a minimum of 15 min. Following slicing, brains were allowed to equilibrate in normal aCSF (in mM: 124 NaCl, 4.4 KCl, 2 CaCl_2_, 1.2 MgSO_4_, 1 NaH_2_PO_4_, 10 glucose, 26 NaHCO_3_, 34° C) for at least 30 minutes. Next, slices were transferred to the recording chamber and allowed to equilibrate in oxygenated aCSF (28-30 °C) perfused at 2 mL/min for an additional 30 minutes. Recordings examining cell excitability were performed in current clamp using K-gluconate intracellular recording solution (K-gluconate 135, NaCl 5, MgCl_2_ 2, HEPES 10, EGTA 0.6, Na_2_ATP 4, Na_2_GTP 0.4). Recordings examining synaptic currents were performed with either in CsCl intracellular solution (130 CsCl, 1 EGTA, 10 HEPES, 2 ATP, 0.2 GTP) or Cs-Methanosulfonate (in mM: 117 Cs methanesulfonic acid, 20 HEPES, 0.4 EGTA, 2.8 NaCl, 5 TEA, 2 ATP, 0.2 GTP) intracellular solutions. CsCl recordings were conducted in kynurenic acid (3mM) to block glutamatergic currents. *Ex vivo* ChR2 stimulation for whole-cell recording was performed using an 470 nM LED from Thor Labs or CoolLED.

### Blood Ethanol Content

Blood ethanol content (BEC) was measured by administering a dose of 2.0 g/kg (20% ethanol w/v, i.p.). Mice were restrained (<2 min) in plexiglass tubes (Braintree Scientific, Braintree, MA) and a scalpel was used to make a small nick in the mouse tail. Blood was collected in a heparinized capillary tube at 30 and 60 minutes following the injection. The plasma was removed and analyzed for BEC using an Analox-G-5 analyzer (Analox Instruments, Lunenbug, MA).

### Homecage Drinking Paradigms

*2-bottle choice* In their homecage, mice were given 24 hour access to a bottle of containing a bottle of test fluid and a bottle of water. The concentration of the test fluid escalated over the course of the experiment at 3 days/dose. These solutions were ethanol (3, 6, 10% w/v, unsweetened), sucrose (0.1, 0.3, 1, 2, 3% w/v), saccharin (0.003, 0.001, 0.03, 0.1% w/v), and quinine (1, 3, 10, 30, 100, 300 μM). We weighed the bottles every 24 hours and switched the side of the cage where the test bottle was located daily. We report these data as the average drinking values for each mouse averaged over the course of the 3 days.

*Intermittent Access (IA)* was performed as described by Hwa *et al*. (2011). Briefly, mice had access to both a bottle of 20% (w/v) ethanol (unsweetened), and water in their homecage on Monday, Wednesday, and Friday. On other days, they only had access to 2 bottles of water. Bottles were rotated with each exposure to ensure that animals did not associate ethanol or water with a particular side of the cage.

### Locomotor and Anxiety Assays

All locomotor and anxiety assays were performed using Ethovision XT tracking software (Noldus Information Technology, Netherlands) to measure location, distance moved, and velocity.

*RTPP* Mice were placed in an apparatus (50 x 50 x 25 cm) that was divided down the middle with a door for exploration on both sides, and which had no distinguishing features on either side. For 20 minutes, mice were allowed to explore the apparatus and received optical stimulation (20 Hz for the ChR2 animals, and constant stimulation for the IC++ animals, 473 nm, 10 mW, Arduino UNO, or Master 8, AMP Instruments, Israel) on one side (counterbalanced) and no stimulation on the other side.

*oICSS* First cohort: *NTS*^CeA➔PBN^::ChR2 (n=14) and control (n=11) mice were food-restricted to 80% of their normal food intake for 2 days before optical intracranial self-stimulation (oICSS). They were tethered to the laser and placed in the chamber (15.9 cm x 14.0 cm x 12.7 cm; MedAssociates, VT, USA) for 1 hour. Both nose ports (active and inactive) were baited with a very small amount of their normal feed to encourage exploration. A dim house light flashed when the animal poked the active port along with 5 seconds of stimulation during which time further pokes had no effect (20 Hz or 40 Hz, 473 nm, 10 mW). Second cohort: *NTS*^CeA➔PBN^::ChR2 (n=8) and control (n=7) mice were not food restricted and ports were baited with a small amount of Froot Loops^TM^ (Kellogg’s). Mice that were fed *ad libitum* did not exhibit reduced motivation to poke for stimulation therefore we collapsed the data across cohorts.

*Open field.* Mice were allowed to explore the open field (50 x 50 cm) for 30 minutes where distance traveled, and velocity were measured (Ethovision, Noldus, Amsterdam).

*Light-dark box.* Mice were placed into the dark enclosed side of the apparatus (Med Associates) and time spent in the light side and entries to the light were monitored for 15 minutes (Ethovision, Noldus, Amsterdam).

*Elevated Plus Maze.* Mice were placed in the center of the apparatus at the beginning of the test. CeA*^NTS^*::casp and control mice were given 5 minutes to explore the open arm, closed arm, and center portion of the maze, and time spent in arms, center, and number of entries were monitored. *NTS*^CeA➔PBN^::ChR2 and control mice were similarly monitored but given 5 minutes to explore the maze without stimulation, 5 minutes with stimulation (20 Hz, 473 nm, 10 mW) and an additional 5 minutes without stimulation (Ethovision, Noldus, Amsterdam).

*Marble burying.* 12 marbles were placed on a 5 cm deep layer of corncob bedding in a standard size mouse cage (39x20x16 cm) in a grid-like fashion. Mice were then placed in the cage for 30 minutes and the degree of marble burying was hand-scored. If a marble was more than ½ way buried it was considered buried. The experimenter was blinded to the viral treatment group prior to the experiment.

*Novelty-suppressed feeding.* Mice were singly-housed a week prior to testing. 48 hours prior to testing, animals were allowed to consume a Froot Loop^TM^ in their homecage. Food was then removed from the homecage for 24 hours. Mice were then placed in a corner of an open field (26.7x48.3 cm) at the center of which we placed a single Froot Loop^TM^ on filter paper. Latency to feed was measured as the time required for the mouse to begin to consume the Froot Loop^TM^. If the mouse had not approached the fruit loop after 10 min, it was removed from the open field and scored as 10 min. Immediately following, the mouse was returned to its homecage and allowed to freely consume Froot Loop^TM^ for 10 min. If the mouse did not consume any Froot Loop^TM^ in the homecage, it was not included for this measurement.

### Optical stimulation consumption paradigm

Mice were habituated to Ethovision Phenotyper boxes (Noldus) over the course of 4 days for 3 hours each. Mice were tethered to the optical commutator, and had access to a bottle of the test fluid and normal chow throughout the habituation period. Over the subsequent 4 days, mice were placed in the same boxes, again with their standard mouse chow and the test fluid in a bottle with a Lick-O-Meter (Noldus) attached. The mice received either optical stimulation across 3 hours (473 nm, 20 Hz, 10mW, 5 min on-off cycles, Fig 9A), or no stimulation (counterbalanced) for within animal comparison (repeated measures two-way ANOVA). Stimulation was delivered in a non-contingent fashion, in order to avoid pairing any particular part of the chamber with the stimulation and producing an RTPP-like effect as seen in Figure 8D. The test fluids were water, 6% (w/v) ethanol, 1% (w/v) sucrose, 0.003% (w/v) saccharine, and 100 μM quinine.

### Statistical Analysis

Data are presented as mean ± SEM. Significance is presented as *p<0.05, **p<0.01, ***p<0.001, ****p<0.0001. All statistical analyses were performed using GraphPad Prism version 6.02 for Windows, GraphPad Software, La Jolla California USA, www.graphpad.com. For the *Fos/Nts in situ* experiment, comparisons were planned between the ethanol and water groups based on the results from the experiments in the caspase drinking studies. Following that, we performed one-way ANOVAs with *Dunnett’s* post-hoc tests (referred to as Dunn’s post-hoc test in Prism) using the water group as the control group. In the caspase experiments we used a Student’s t-test. Optogenetic behavioral data was subjected to a matched 2-way ANOVA were applicable, followed by post-hoc Bonferroni-corrected t-tests if a significant interaction was detected. Where ANOVAs were not applicable, the data was subjected to a Student’s t-test. Data are reported as the mean ± SEM. The fluid consumption values for the FISH experiment were reported as standard deviation (SD) to convey variability in the drinking.

One *NTS*^CeA^::eYFP (control) animal was removed from the caspase drinking studies due to extremely low ethanol consumption. It consumed no more than 2.1 g/kg ethanol average per week and its preference for ethanol was greater than 2 standard deviations from the mean for control animals. One *NTS*^CeA->PBN^::ChR2 was removed from the water-drinking phenotyper experiment. Stimulation-day drinking for this mouse was a ROUT outlier from <u>all</u> other water drinking days (stim and non-stim, *NTS*^CeA->PBN^::ChR2 and *NTS*^CeA->PBN^::eYFP).

## Results

### NTS neurons in the CeA express a variety of markers

We first explored how *Nts*-expressing neurons overlap with other previously described genetically-defined populations in the central amygdala (CeA). Using dual fluorescent *in situ* hybridization (FISH) across the entire CeA, we examined neuronal overlap with cells expressing mRNA for corticotropin releasing hormone (also known as corticotropin releasing factor, *Crh*), corticotropin-releasing hormone receptor 1 (also known as CRF receptor 1, *Crhr1*), preprodynorphin (*Pdyn*), protein kinase c delta (*Pkcδ)*, and somatostatin (*Sst*). We found that CeA *Nts*-expressing neurons largely express *Crh* and *Crh1* (Fig 1). Surprisingly, we found that a third of CeA *Nts* neurons express *Pkcδ*, a population that has been reported to have limited overlap with CeA *Crf* cells (Cai et al., 2014). One third of *Nts* CeA neurons express *Sst,* a population that has been implicated in the switch between passive and active stress coping mechanisms (Yu et al., 2016). Lastly, about two-thirds of CeA-NTS labeled neurons also express *Pdyn*, the precursor of the endogenous ligand for the kappa opioid receptor, dynorphin (Chavkin et al., 1982).

**Fig 1:**
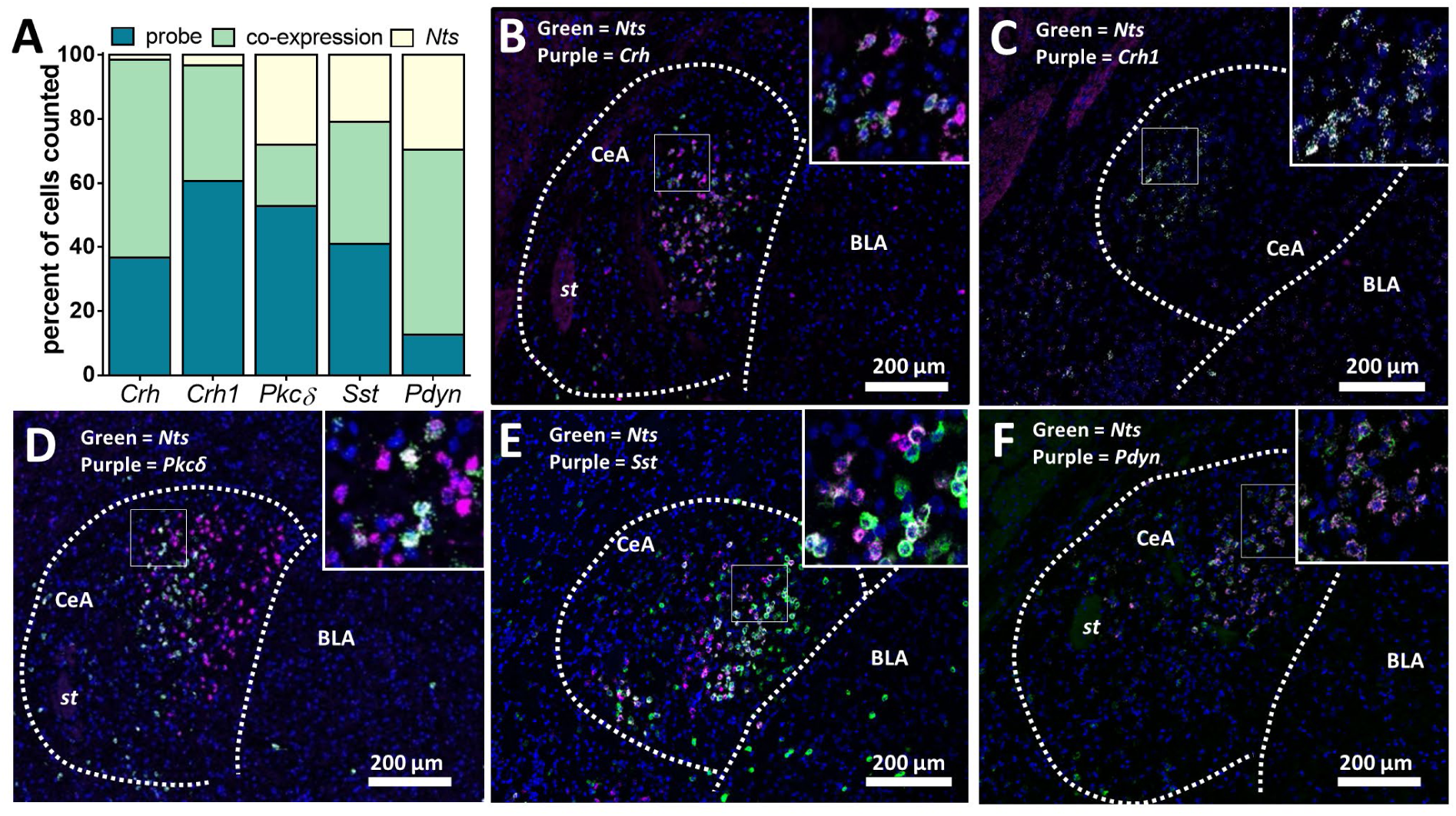
Nts neurons in the CeA express a variety of markers. (a) Quantification of dual FISH in the CeA for *Nts* co-localization with *Crh*, *Crh1*, *Pkcδ*, *Sst*, and *Pdyn*. (**b-f**) Representative confocal images with *Nts* (green), probe (purple), and DAPI (blue). (b) 98% of *Nts* neurons expressed *Crh*, and 37% of *Crh* expressed *Nts* (n=3 mice, 5-6 slices/mouse). (c) 92% of *Nts* neurons expressed *Crh1*, and 63% of *Crh1* expressed *Nts* (n=4 mice, 5-6 slices/mouse). (d) 41% of *Nts* expressed *Pkcδ* and 27% of *Pkcδ* neurons expressing *Nts* (n=4 mice, 2-4 slices/mouse). (e) 65% of *Nts* expressed *Sst* and 48% of *Sst* neurons expressing *Nts* (n=4 mice, 2-4 slices/mouse). (f) 48% of *Nts* expressed *Pdyn* and 82% of *Pdyn* neurons expressed *Nts* (n=4 mice, 5-6 slices/mouse). (Green= *Nts*, Purple= probe, Blue= DAPI, st= stria terminalis, CeA= central amygdala, BLA = basolateral amygdala, all scale bars 200 µm).

### Ablation of NTS^CeA^ neurons decreases ethanol consumption in two-bottle choice

To determine if *NTS*^CeA^ neurons play a role in ethanol-related behavior, we used NTS-IRES-Cre-recombinase (NTS-Cre) mice (Leinninger et al., 2011) in conjunction with viral manipulations in the CeA.

First, we validated the fidelity and penetrance of *Cre* in the CeA of this line. Using FISH (Fig 2A), we double-labeled *Nts* and *Cre* mRNA in CeA slices from 5 separate NTS-Cre mice. We found that 61.4% of *Nts* mRNA-expressing cells also expressed *Cre* and we found that 82.2% of *Cre* mRNA-expressing cells also expressed *Nts* mRNA. These data indicate this is a high-fidelity Cre line with strong penetrance.

**Fig 2:**
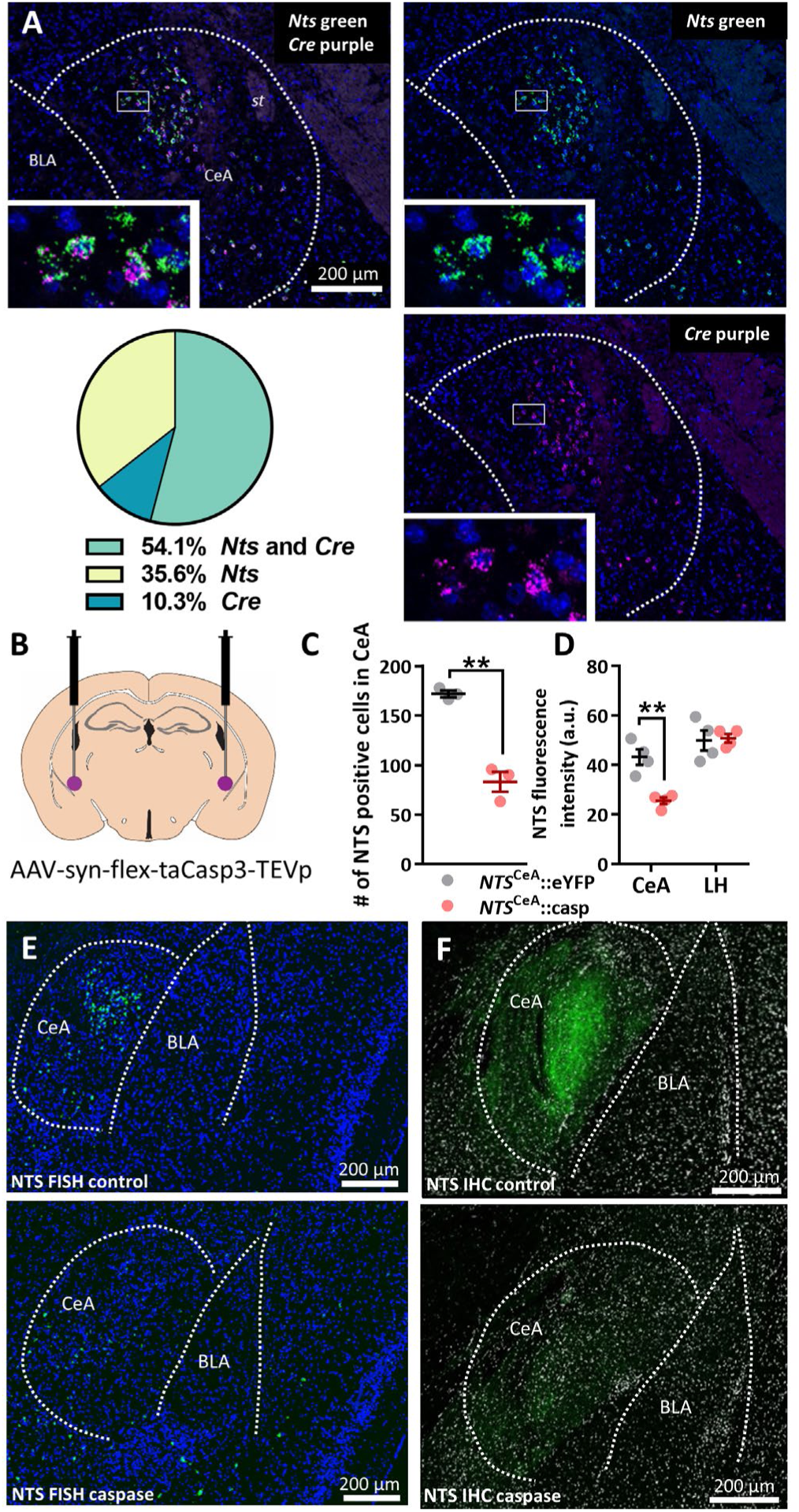
NTS-Cre line and caspase manipulation validation. (a) Dual FISH of *Nts* (green) and *Cre* (purple) in the CeA with DAPI (blue). 61.4% of *Nts* mRNA-expressing cells (241.2 ± 29.7 *Nts*+ cells per slice) also expressed *Cre* (145.4 ± 23.7 *Nts*+*Cre*+ cells per slice) and 82.2% of *Cre* mRNA-expressing cells (173.2 ± 22.8 *Cre*+ cells per slice) also expressed *Nts* mRNA (n=3 mice, 5-6 slices/mouse). (**b**) Diagram of CeA injection site. (**c**) Quantification of cells FISH labeled for *Nts* in the CeA from *NTS*^CeA^::casp (n=3) and *NTS*^CeA^::eYFP animals (n=3, unpaired t-test: t(4)=8.425, p=0.0011). (**d**) Caspase ablation decreased NTS immunoreactivity as measured in arbitrary units (a.u.) in the CeA (unpaired t-test: t(6)=5.090, p=0.0022), but not in the LH (unpaired t-test: t(6)=0.1956, p=0.8514). Representative images of *in situ* (**e**) and IHC (**f**). **p<0.01 unpaired t-test.

We next injected a Cre-dependent virus encoding a modified pro-caspase 3 and TEV protease (AAV5-Ef1a-FLEX-taCasp-TEVp; Yang et al., 2013) into the CeA of NTS-Cre mice to selectively lesion *NTS*^CeA^ neurons (*NTS*^CeA^::casp, Fig 2B). This strategy resulted in a 51.7% reduction in NTS-positive cells in the CeA (Fig 2C) and a 40.9% reduction in CeA-NTS immunoreactivity, without altering NTS-ir in the neighboring LH (Fig 2D). Control animals were injected with a Cre-dependent eYFP construct (*NTS*^CeA^::eYFP).

Due to the importance of the CeA in ethanol consumption (Gilpin et al., 2015), we hypothesized the loss of *NTS*^CeA^ neurons would alter voluntary ethanol consumption in a continuous 2-bottle choice paradigm. *NTS*^CeA^::casp mice showed significant decreases in ethanol consumed in 24-hour 2-bottle choice drinking when compared to *NTS*^CeA^::eYFP controls (Fig 3A; Two-way ANOVA: interaction, F_(2,42)_= 6.340, p=0.0039; ethanol concentration, F_(2,42)_=98.23, p<0.0001; ablation, F_(1,21)_=16.52, p=0.0006), with no effect of preference for the ethanol bottle (Fig 3B; Two-way ANOVA: interaction, F_(2,42)_=1.793, p=0.1790; ethanol concentration, F_(2,42)_=7.727, p=0.0014; ablation, F_(1,21)_=3.283, p=0.0843). *NTS*^CeA^::casp animals also showed decreased liquid consumption at lower ethanol concentrations, which was driven by increased total drinking by the *NTS*^CeA^::eYFP mice at lower ethanol concentrations (Fig. 3F; Two-way ANOVA: interaction, F_(2,42)_=6.551, p=0.0033; ethanol concentration, F_(2,42)_=47.02, p<0.0001; ablation, F_(1,21)_=9.208, p=0.0063). Because of this, we next determined whether *NTS*^CeA^::casp mice showed general differences in liquid consumption compared to controls and measured water drinking over 5 days. *NTS*^CeA^::casp mice drank the same amount of water as *NTS*^CeA^::eYFP mice (Fig 3G; Two-way ANOVA: interaction, F_(4,44)_=2.459, p=0.0593; ablation, F_(1,11)_=1.005, p=0.3377; day, F_(4,44)_=2.714, p=0.0418), confirming that *NTS*^CeA^ ablation affects ethanol consumption as opposed to general liquid consumption.

**Fig 3:**
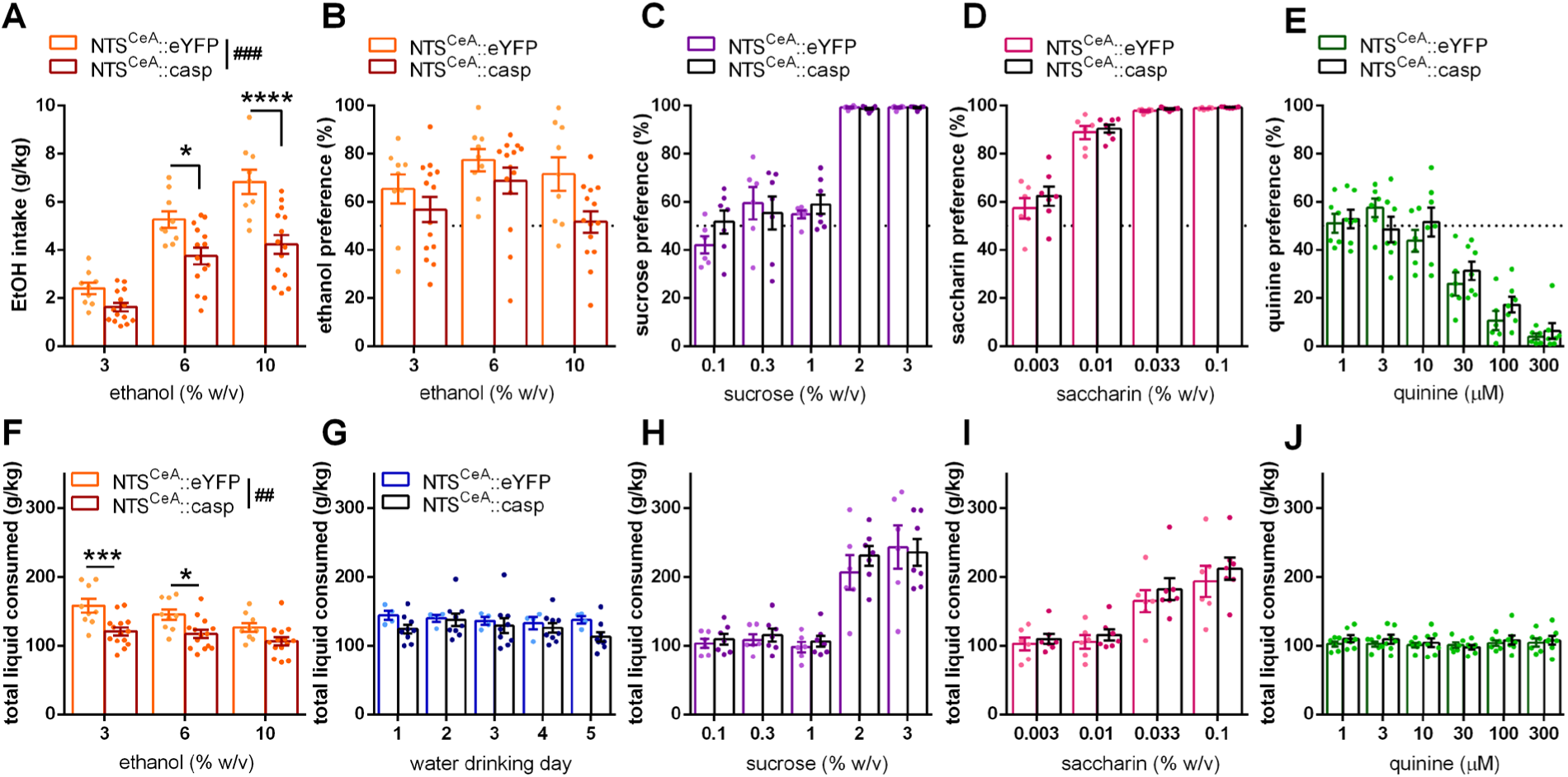
Ablation of NTS neurons in the CeA decreases ethanol drinking in 2-bottle choice. (a) *NTS*^CeA^::casp mice (n=14) drank significantly less ethanol than *NTS*^CeA^::eYFP control animals (n=9). (b) Preference for the tastant bottle was not significantly different between these groups for either ethanol, (**c)** sucrose (eYFP n=6, casp n=7), (**d**) saccharin (eYFP n=6, casp n=7) or (**e**) quinine (eYFP n=6, casp n=7). (**f**) Liquid consumed was significantly different between *NTS*^CeA^::casp and *NTS*^CeA^::eYFP groups when the mice consumed ethanol, but not when they consumed (**g**) water (eYFP n=4, casp n=9), (**h**) sucrose, (**i**) saccharin, or (**j**) quinine. Bonferroni-corrected t-tests: *p<0.05, ***p<0.001, ****p<0.0001. ANOVA main effects: ^##^p<0.01 ^###^p<0.001.

In order to determine whether this decrease in alcohol consumption was due to an increase in aversion to a bitter tastant, or decreased hedonic value for a rewarding fluid, we performed a series of two-bottle choice preference tests with multiple caloric and non-caloric tastants. In a new cohort of animals, the *NTS*^CeA^::eYFP and *NTS*^CeA^::casp groups showed no difference in preference for sucrose (Fig 3C; Two-way ANOVA: interaction, F_(4,44)_=0.8346, p=0.5106; concentration, F_(4,44)_=76.89, p<0.0001; ablation, F_(1,11)_=0.8047, p=0.3889), saccharin (Fig 3D; Two-way ANOVA: interaction, F_(3,33)_=0.4399, p=0.7260; concentration, F_(3,33)_=134.0, p<0.0001; ablation, F_(1,11)_=1.063, p=0.3246) or quinine (Fig 3E; Two-way ANOVA: interaction, F_(5,55)_=1.139, p=0.3511; concentration, F_(5,55)_=52.53, p<0.0001; ablation, F_(1,11)_=0.6999, p=0.4206). Additionally, the *NTS*^CeA^::eYFP and *NTS*^CeA^::casp groups did not differ in the consumed volume (liquid g/kg) of any of these tastants (Sucrose Two-way ANOVA: interaction, F_(4,44)_=0.4449, p=0.7755; sucrose concentration, F_(4,44)_=109.1, p<0.0001; ablation, F_(1,11)_=0.2132, p=0.6533); Saccharin Two-way ANOVA: interaction, F_(3,33)_=0.2004, p=0.8954; saccharin concentration, F_(3,33)_=126.2, p<0.0001; ablation, F_(1,11)_=8.016, p=0.3781); Quinine Two-way ANOVA: interaction, F_(5,55)_=0.7687, p=0.5764; quinine concentration, F_(5,55)_=52.51, p<0.0001; ablation, F_(1,11)_=1.254, p=0.2866). Lastly, the daily total liquid consumed was not different between the *NTS*^CeA^::eYFP and *NTS*^CeA^::casp groups for either sucrose (Fig 3H; Two-way ANOVA: interaction, F_(4,44)_=0.4976, p=0.7375; concentration, F_(4,44)_=69.17, p<0.0001; ablation, F_(1,11)_=0.2049, p=0.6596), saccharin (Fig 3I; Two-way ANOVA: interaction, F_(3,33)_=0.2906, p=0.8318; concentration, F_(3,33)_=86.01, p<0.0001; ablation, F_(1,11)_=0.5694, p=0.4664) or quinine (Fig 3J; Two-way ANOVA: interaction, F_(5,55)_=1.092, p=0.3754; concentration, F_(5,55)_=2.456, p=0.0444; ablation, F_(1,11)_=0.2943, p=0.5983). These data suggest that the decrease in ethanol intake measured in *NTS*^CeA^::casp animals was not due to changes in general fluid intake, motivation to drink rewarding fluids in general, or aversion to bitter tastants, but was instead specific for ethanol.

We wanted to verify that genetic ablation of *NTS*^CeA^ neurons did not result in gross changes in body weight or movement. We measured body weight for a month following stereotactic surgery and found that this lesion did not alter body weight (Fig 4A; Two-way ANOVA: interaction, F_(26, 208)_=0.9646; day, F_(26,208)_= 40.11, p<0.0001, p=0.5180; ablation, F_(1,8)_=0.1154, p=0.7428). We also tested the animals in an open field and found no changes in locomotor behavior measured as either distance travelled (Fig 4B; Two-way ANOVA: interaction, F_(2,36)_=0.9989, p=0.3783; time, F_(2,36)_=109.3, p<0.0001; ablation, F_(1,18)_=0.1886, p=0.6693) or velocity (Fig 4C; Two-way ANOVA: interaction, F_(2,38)_=0.9970, p=0.3784; time, F_(2,38)_=98.55, p<0.0001; ablation, F_(1,19)_=0.2698, p=0.6095). We next wanted to verify that *NTS*^CeA^::casp animals did not have differences in other ethanol-related traits that might be responsible for their blunted drinking, specifically sedation following a high dose of ethanol and ethanol metabolism. *NTS*^CeA^ neuron ablation did not change sedation in response to ethanol (Fig 4D; 3.2 g/kg dose: Unpaired t-test t(10)=0.0001, p=0.9999; 4.5 g/kg dose: Unpaired t-test t(11)=0.5696, p=0.5804) or ethanol metabolism as measured by blood ethanol content following an i.p. injection of 2.0 g/kg of ethanol (Fig 4E; Two-way ANOVA: interaction, F_(1,8)_=1.270, p=0.2924; time, F_(1,8)_=1.964, p=0.1987; ablation, F_(8,8)_=2.538, p=0.1046).

**Fig 4:**
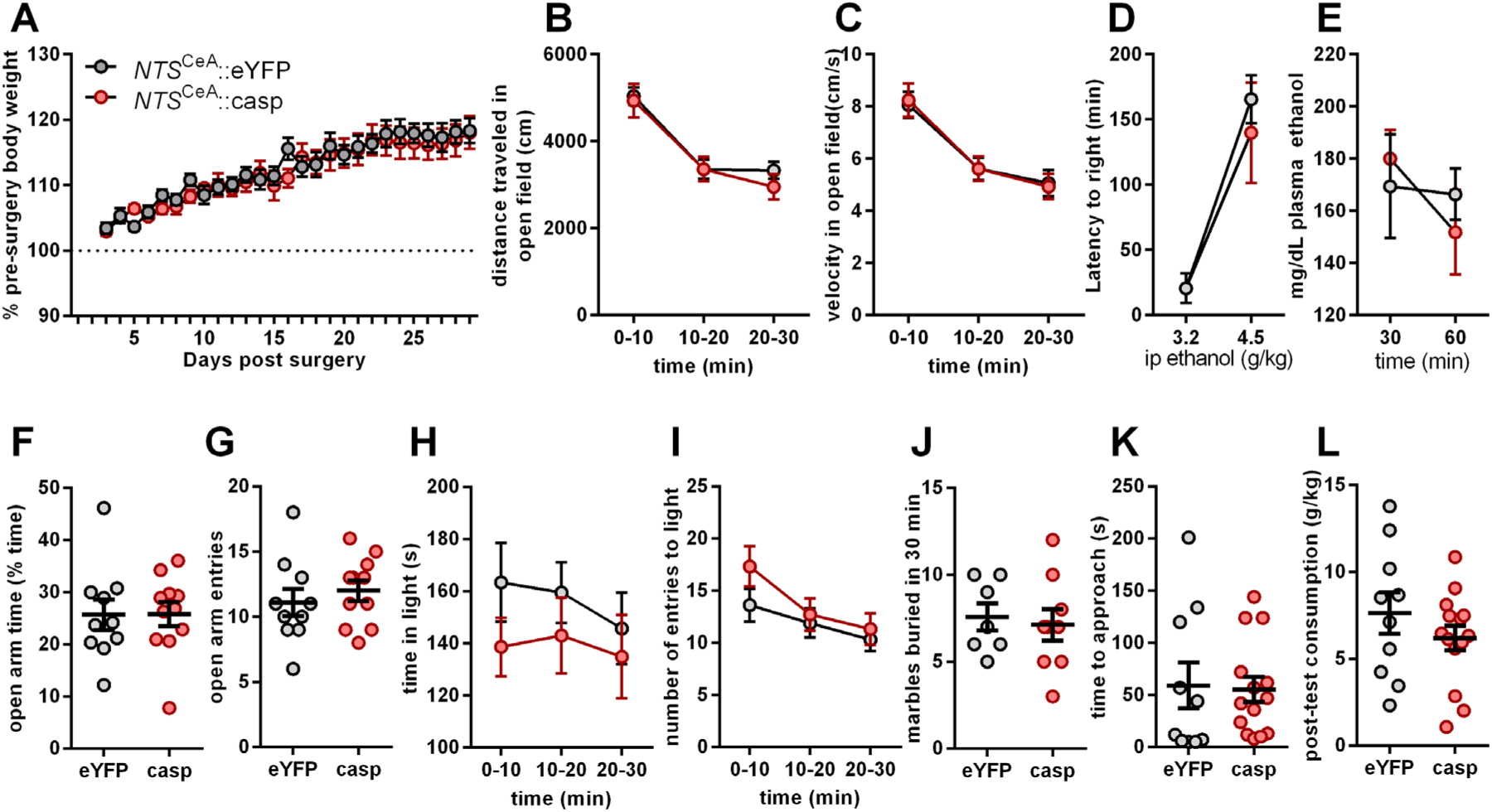
Ablation of NTS neurons in the CeA does not alter ethanol metabolism, body weight or anxiety-like behavior. (**a**) *NTS*^CeA^::casp mice (n=5) and *NTS*^CeA^::eYFP mice (n=5) had similar growth curves post-surgery. (**b**) *NTS*^CeA^ ablation did not affect either distance traveled or (**c**) velocity in an open field (eYFP n=9, casp n=11). (**d**) *NTS*^CeA^ ablation did not affect latency to right following a 3.2 g/kg or 4.5 g/kg ethanol i.p. injection (eYFP n=6, casp n=7). (**e**) Blood alcohol concentrations (BACs) following administration of 2.0 g/kg i.p. ethanol was not affected by *NTS*^CeA^ ablation (eYFP n=5, casp n=5). (**f**) *NTS*^CeA^ ablation did not affect either time spent in or (**g**) entries to the open arms of an elevated plus maze (eYFP n=10, casp n=11). (**h**) *NTS*^CeA^ ablation did not affect either time spent in or (**i**) entries to the light side of a light-dark box (eYFP n=16, casp n=18). (**j**) *NTS*^CeA^::casp mice (n=9) and *NTS*^CeA^::eYFP mice (n=7) buried similar numbers of marbles in a marble-burying test. (**k**) *NTS*^CeA^::casp mice (n=14) and *NTS*^CeA^::eYFP mice (n=10) were not different in time to approach the food in the novelty-suppressed feeding task or in (**l**) the 10 minute consumption post-test.

### Ablation of NTS^CeA^ neurons does not impact anxiety-like behavior

Given the potential role of the CeA in anxiety, we also conducted a series of behavioral tests to measure anxiety-like responses. Genetic ablation failed to alter anxiety-like behaviors as measured by: time spent in and entries to the open arms of an elevated plus maze (Fig 4F-G; *time spent*: Unpaired t-test: t(19)=0.03167, p=0.9751; *entries*: Unpaired t-test: t(19)=0.6992, p=0.4929), time spent in and entries to the light side of a light-dark box (Fig 4H-I; *time spent*: Two-way ANOVA: interaction, F_(2,64)_=0.3707, p=0.6917; time, F_(2,64)_=1.203, p=0.3071; ablation, F_(1,32)_=1.000, p=0.3247; *entries*: Two-way ANOVA: interaction, F_(2,60)_=1.452, p=0.2422; time, F_(2,60)_=14.63, p<0.0001; ablation, F_(1,30)_=0.7529, p=0.3924), marble-burying (Fig 4J; Unpaired t-test: t(14)=0.3716, p=0.7158) or novelty-suppression of feeding (Fig 4K-L; Unpaired t-test: t(22)=0.1597, p=0.8746). Based on these data, genetic ablation of *NTS*^CeA^ neurons selectively reduced alcohol consumption without affecting motor function, the sedative-hypnotic effects of ethanol, blood ethanol clearance, or anxiety-like behavior.

### Ablation of NTS^CeA^ neurons decreases ethanol consumption in Intermittent Access

Because of the ethanol dose effect observed with our initial 2-bottle choice experiments (Fig 3A), we next examined whether ablation of *NTS*^CeA^ neurons would alter ethanol consumption in a drinking paradigm with a longer schedule of access and a higher dose of alcohol. We used an intermittent access (IA) drinking paradigm in an attempt to increase alcohol consumption. *NTS*^CeA^::casp mice again showed significant decreases in ethanol consumed across all weeks as compared to *NTS*^CeA^::eYFP controls (Fig 5A; Two-way ANOVA: interaction, F_(6,126)_=0.4321, p=0.8564; week, F_(6,126)_=2.539, p=0.0235; ablation, F_(1,21)_=11.19, p=0.0031) as well as cumulative ethanol consumption (Fig 5B; Two-way ANOVA: interaction, F_(20,380)_=13.53, p<0.0001; day, F_(20,380)_= 194.5, p<0.0001; ablation, F_(1,19)_= 11.69, p=0.0029. Bonferroni-corrected post-hoc tests show significant difference between *NTS*^CeA^::casp and *NTS*^CeA^::eYFP at days 26 through 47). Total liquid consumed was unaffected whether measured by week (Fig 5C; Two-way ANOVA: interaction, F_(6,126)_=1.525, p=0.1752; week, F_(6,126)_=8.358, p<0.0001; ablation, F_(1,21)_=0.00005215, p=0.9943) or cumulative intake (Fig 5D; Two-way ANOVA: interaction, F_(20,420)_=0.1298, p>0.9999; day, F_(20,420)_=861.7, p<0.0001; ablation, F_(1,21)_=0.01703, p=0.8976). *NTS*^CeA^::casp mice also showed a significant decrease in preference for the ethanol bottle (Fig 5E; Two-way ANOVA: interaction, F_(6,126)_=0.7778, p=0.588; week, F_(6,126)_=3.992, p=0.0011; ablation, F_(1,21)_=15.88, p=0.0007). Lastly, we compared the total amount consumed at the end of the 7 weeks of IA. *NTS*^CeA^::casp mice consumed significantly less total ethanol than *NTS*^CeA^::eYFP mice (Fig 5F; Unpaired t-test t(21)=3.413, p=0.0026), with no detectable difference in total liquid consumed (Fig 5G; Unpaired t-test: t(21)=0.04085, p=0.9678). These experiments suggest that *NTS*^CeA^ neurons regulate ethanol consumption across multiple dose ranges and schedules of access.

**Fig 5:**
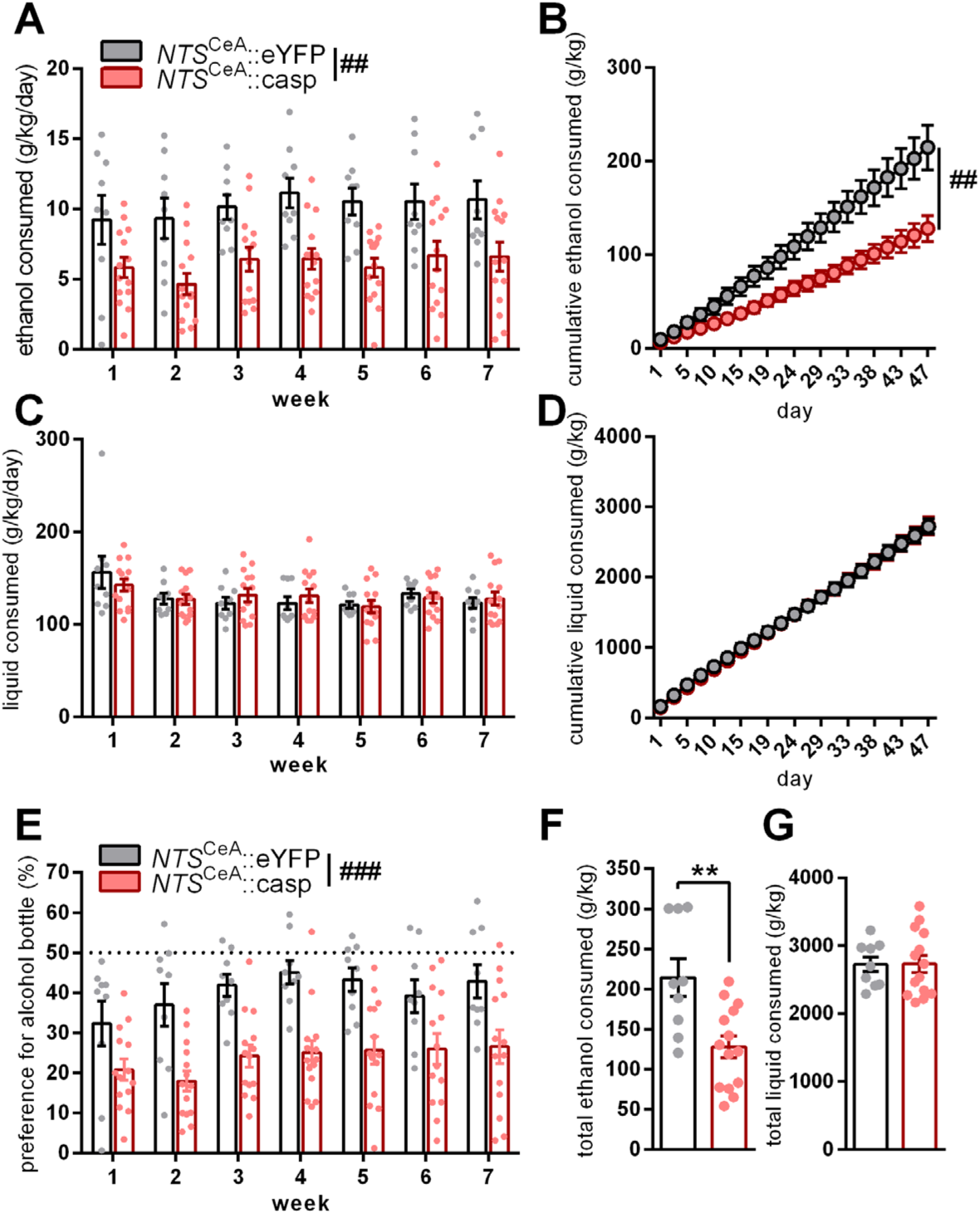
Ablation of NTS neurons in the CeA decreases ethanol drinking and preference in an intermittent access (IA) paradigm. (**a**) *NTS*^CeA^::casp mice (n=14) consume less ethanol than *NTS*^CeA^::eYFP mice (n=9) in an IA paradigm whether measured weekly or (**b**) cumulatively. (**c**) General liquid consumption was not affected by caspase ablation whether measured by week or (**d**) cumulatively. (**b**, **d**) Days are numbered from the beginning of the experiment (each circle represents an ethanol drinking day). (**e**) Preference for the ethanol bottle was significantly different between the *NTS*^CeA^::casp and *NTS*^CeA^::eYFP mice. (**f**) Cumulative ethanol consumption over all 7 weeks of IA was significantly different between the *NTS*^CeA^::casp and *NTS*^CeA^::eYFP mice, but cumulative liquid consumption over the same period was not (**g**). Unpaired t-tests: **p<0.01. ANOVA main effects: ^##^p<0.01 ^###^p<0.001.

### Neurons in the central amygdala are activated by various tastants

In order to determine whether *Nts* neurons in the CeA would be activated following voluntary consumption of ethanol, we performed dual fluorescence *in situ* hybridization (FISH) for *Nts* and *Fos* in CeA slices. Singly-housed male C57BL/6J mice were allowed access to either water, 6% ethanol, 1% sucrose, 0.03% saccharin, or 100 μM quinine and for 2 hours during 4 consecutive days. On the 5^th^ day, the mice consumed fluid for 1 hour and were euthanized 30 minutes later for FISH. The average fluid consumption for these groups was 8.34 g/kg (4.49 SD) for water, 10.44 g/kg (6.18 SD) for ethanol, 32.84 g/kg (15.96 SD) for sucrose, 36.25 g/kg (8.86 SD) for saccharin, and 5.34 g/kg (3.94 SD) for quinine. This homecage drinking failed to induce changes in *Fos* mRNA expression in the CeA when analyzed in total (Fig 6A), however, work investigating genetically-defined subpopulations of neurons in the CeA suggests that *Nts* neurons can be subdivided into functionally separate medial (CeA_M_) and lateral (CeA_L_) populations (Kim et al., 2017). We thus subdivided the images into CeA_M_ and CeA_L_, focusing on slices located from - 1.1 to -1.8 posterior to Bregma, where it was easier to delineate between these two regions. Tastant consumption did not change *Fos* expression when compared to the water group (Fig 6B-C), with the exception of sucrose consumption increasing *Fos* specifically in the CeA_M_ (Fig 6B; Dunnett’s Multiple comparison’s test: water vs sucrose, adjusted p=0.0367). We then examined activation of *Nts* neurons specifically (Fig 6D-F). We performed an *a priori* planned comparison between the water and ethanol groups as the *NTS*^CeA^::casp animals only showed a phenotype for ethanol drinking. Interestingly, ethanol consumption resulted in an increase in the percent of *Fos*-expressing *Nts* neurons in the CeA_L_ (Fig 6F; Unpaired t-test with Welch’s correction: t(9.685)=2.248, p=0.0491). These data suggest that the CeA_L_ group of NTS neurons might be responsible for the ethanol phenotype seen in the *NTS*^CeA^::casp animals.

**Fig 6:**
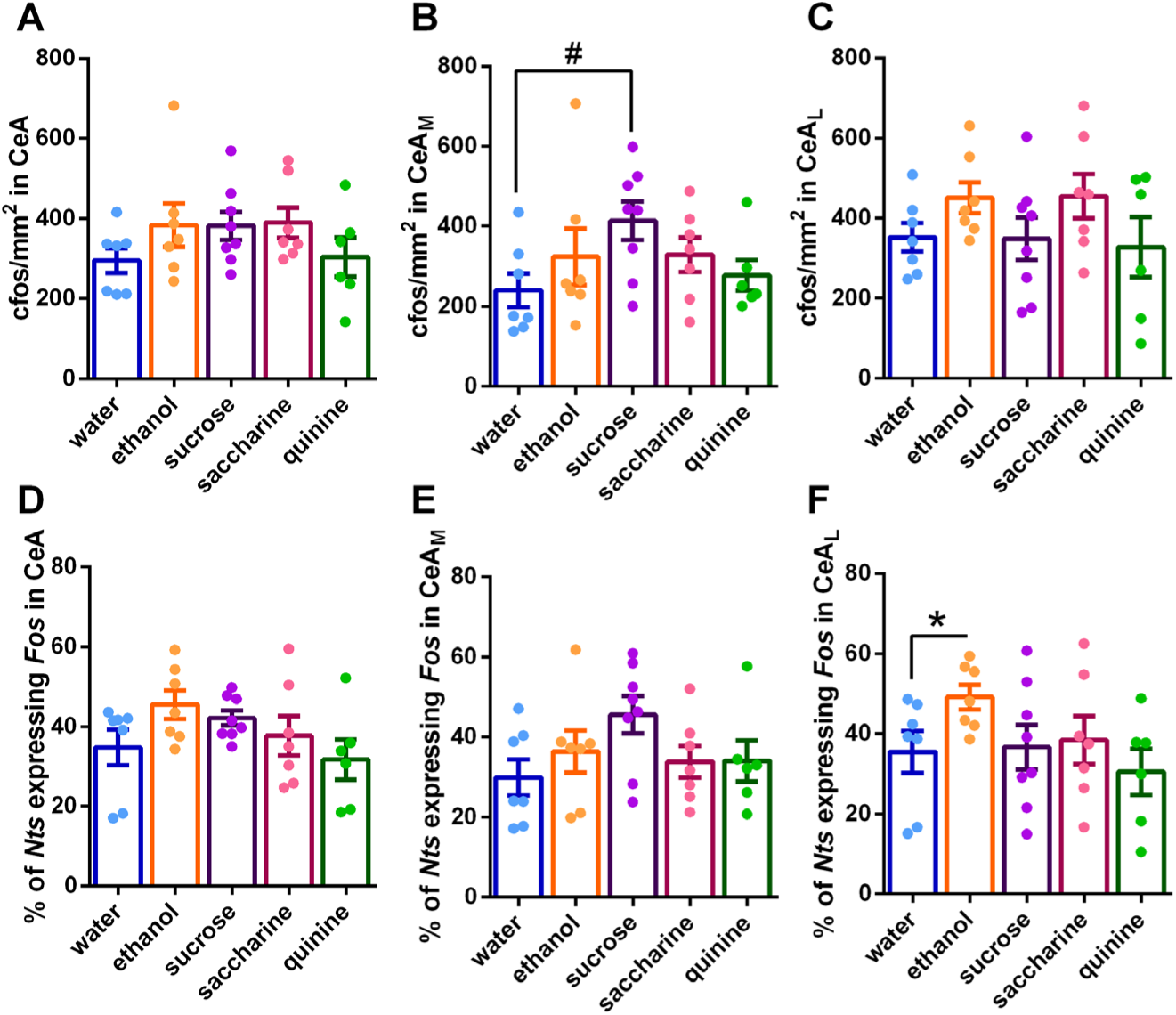
Nts+ neurons in the lateral CeA are activated by ethanol in vivo. C57BL/6J mice consumed either water (n=7), 6% ethanol (n=7), 1% sucrose (n=8), 0.03% saccharin (n=7), or 100 μM quinine (n=6). (**a**) *Fos* expression in the CeA_total_ as a whole was unchanged across all tastants. (**b**) Sucrose consumption increased *Fos* expression in the CeA_M_ but not in (**c**) the CeA_L_. (**d**) The percent of *Nts* neurons expressing *Fos* was unchanged by tastant exposure in the CeA_total_ and (e) CeA_M_. (**f**) Ethanol consumption increased *Fos* expression in *Nts* neurons in the CeA_L_. Planned unpaired t-test: *p<0.05; Dunnetts’s Multiple comparisons test: #p<0.01.

### NTS^CeA^ neurons send a dense projection to the parabrachial nucleus (PBN)

To begin to examine the targets of *NTS*^CeA^ neurons, we injected a Cre-dependent virus expressing channelrhodopsin-2 tagged with eYFP (ChR2-eYFP) into the CeA of NTS-IRES-Cre mice (Fig 7A-B). Using whole-cell *ex vivo* slice electrophysiology and recording in current clamp, we found that 473 nm light stimulation (20 Hz, 5 ms pulse) readily evoked action potentials in *NTS*^CeA^::ChR2 neurons (data not shown). We observed a projection from *NTS*^CeA^ neurons to the hindbrain near the 4^th^ ventricle with robust fluorescence expression in the PBN and the lateral edge of the locus coeruleus (LC, Fig 7C), as well as a projection to the bed nucleus of the stria terminalis (BNST) which was particularly dense in the ventral fusiform subnucleus (Fig 7D). We found significantly greater fluorescence expression in the PBN versus the LC (Fig E; Unpaired t-test: t(6)=14.59, p<0.0001). However, LC neurons extend long dendritic processes into the boundaries of the PBN (Swanson, 1976) so we next sought to determine where *NTS*^CeA^ neurons make functional synaptic connections using electrophysiology.

**Fig 7:**
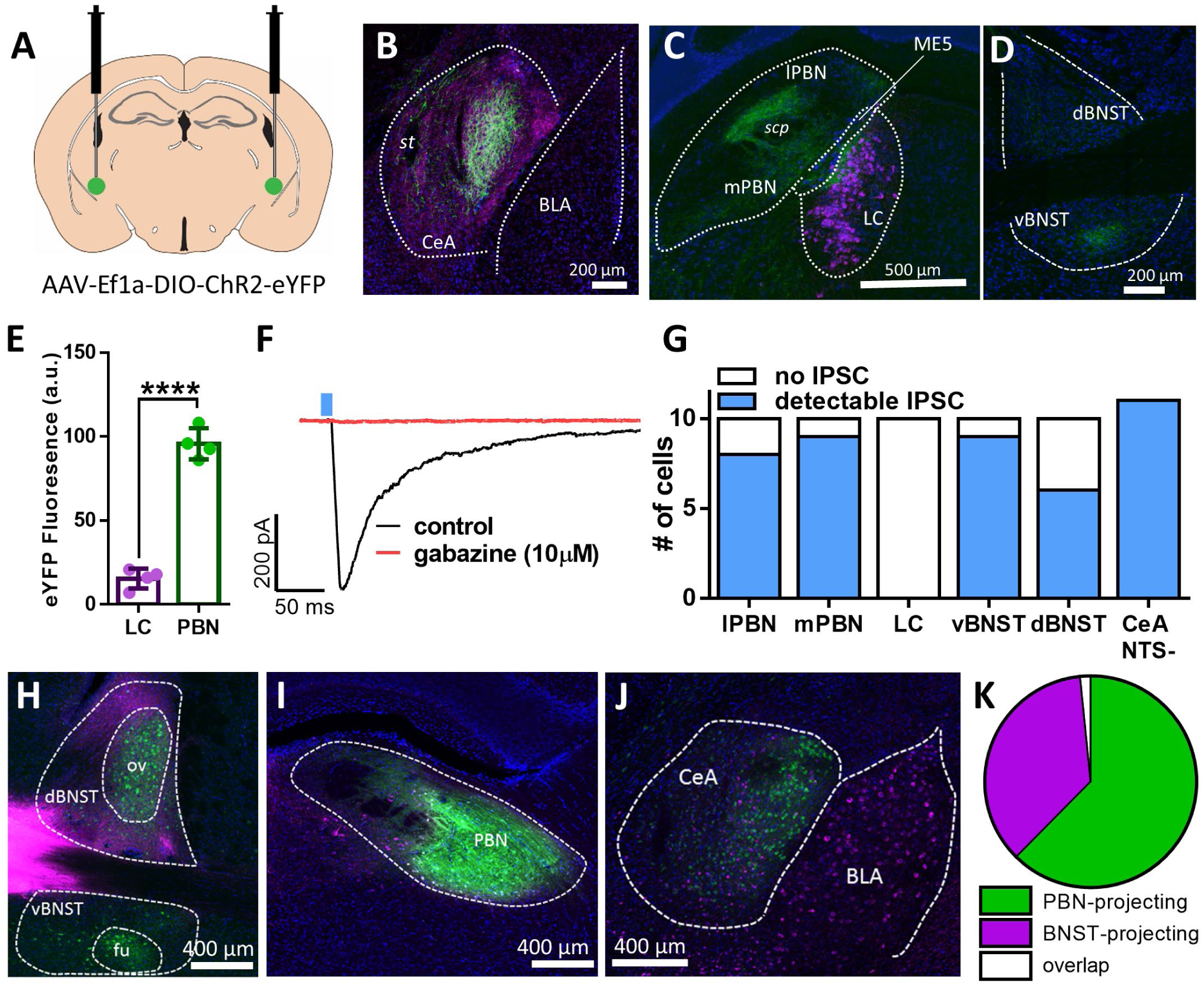
NTS^CeA^ neurons project to the parabrachial nucleus (PBN). (**a**) Diagram of injection site in the CeA of AAV-EF1α-DIO-ChR2-eYFP in the CeA of NTS-IRES-Cre mice. (**b**) Representative image of CeA expression of ChR2-eYFP (green), NTS IHC (purple), and DAPI (blue) in the CeA (*st*= *stria terminalis,* BLA = basolateral amygdala). (**c**) Representative image of hindbrain, *NTS*^CeA^::ChR2-eYFP fibers (green), tyrosine hydroxylase (TH, purple), neurons (blue). (lPBN = lateral parabrachial nucleus, mPBM = medial parabrachial nucleus, LC = locus coeruleus, ME5 = midbrain trigeminal nucleus, *scp* = superior cerebellar peduncle) (**d**) Representative image of expression of *NTS*^CeA^::ChR2-eYFP fibers (green) in the BNST with DAPI staining (blue, dBNST = dorsal portion of the bed nucleus of the stria terminalis, vBNST = ventral portion of the bed nucleus of the stria terminalis). (**e**) PBN has significantly greater eYFP fluorescence intensity (a.u.) as compared to the LC in *NTS*^CeA➔PBN^::ChR2 (n = 4; Unpaired t-test: t(6)=14.59, p<0.0001). (f) Representative trace of oeIPSC in the PBN and its inhibition by gabazine (10 µM). The blue line indicates the delivery of a light pulse (5ms). (**g**) Quantification of cells with light-evoked responses in *NTS*^CeA^ animals in the lPBN (8/10 cells), mPBN (9/10 cells), LC (0/10 cells), vBNST (9/10 cells), dBNST (6/10 cells), as well as eYFP- CeA neurons (11/11). (**h**) Representative BNST image of retrograde cholera toxin-b (CTXb) tracing experiment (ov = oval nucleus of the BNST, fu = fusiform nucleus of the BNST). (**i**) Representative PBN image of retrograde cholera toxin-b (CTXb) tracing experiment. (**j**) Representative CeA image of retrograde cholera toxin-b (CTXb) tracing experiment. Green = cells projecting to the parabrachial nucleus (PBN), purple = cells projecting to the BNST. (**k**) Quantification of cell body fluorescence expression (green and purple CTXb) in the CeA (n = 3 mice). 62.4% of labeled neurons projected to the PBN, 36.0% projected to the BNST, and 1.6% of cells were doubly-labeled.

Monosynaptic input was isolated in whole-cell patch clamp recordings with TTX (500 µM) and 4-AP (1 mM). 473 nm light stimulation (5 ms) of CeA-NTS terminals induced an optically-evoked inhibitory post-synaptic current (oeIPSC) in both the medial and lateral PBN which was blocked by the GABAA receptor antagonist gabazine (10μM; example trace Fig 7F), while no inhibitory or excitatory synaptic currents were observed in the LC (Fig 7G). These data suggest that the *NTS*^CeA^ neurons make functional inhibitory synaptic connections in the lateral and medial portions of the PBN (8 of 10 cells, and 9 of 10 cells respectively) but not the LC (0 of 10 cells, n=6 mice). While we do not know the genetic identity of the PBN neurons receiving this innervation, the possibility remains that these neurons may reciprocally project to the CeA as both *Oxtr*^PBN^ and *Calca*^PBN^ neurons regulate fluid intake (Carter et al., 2013; Ryan et al., 2017). We also verified a synaptic inhibitory *NTS*^CeA^ projection to the BNST which was stronger in the ventral portion (9 of 10 cells) than in the dorsal portion (6 of 10 cells). We also found strong local connections within the CeA. All non-eYFP labeled cells examined (11 of 11 cells, n=4 mice) exhibited an optically evoked IPSC. Interestingly, three of these eYFP- cells were BNST-projecting neurons identified using retrobeads injected into the BNST. This strong local inhibition from *NTS*^CeA^ neurons, in conjunction with our *Fos* FISH tastant study (see above), suggested that cell-body optogenetic stimulation of the entire *NTS*^CeA^ population might not be reflective of the activation of these neurons *in vivo*, thus, we decided to pursue a pathway-specific strategy.

To narrow our focus of target regions, we explored the two nuclei where we observed the densest fiber innervation following the expression of ChR2 in the *NTS*^CeA^ the BNST and PBN. In order to determine whether individual *NTS*^CeA^ neurons collateralize to both the BNST and PBN, we injected the retrograde tracer Alexa-555 cholera toxin-b (CTXb) into the BNST (Fig 7H) and Alexa-488 (CTXb into the PBN (Fig 7I) of the same animal. We found minimal overlap between BNST- and PBN- projecting neurons (1.6%, Fig 7J-K) suggesting that these are distinct cell populations within the CeA. Somewhat surprisingly, we also noted that the BNST- and PBN-projecting neurons in the CeA appear to have a medial-lateral gradient, with the larger population of PBN-projecting neurons located in the CeA_L_. Combining this observation with the significant elevation of *Fos* in the CeA_L_ following moderate ethanol consumption, the established role for the PBN in consummatory behaviors, we hypothesized that the CeA-NTS projection to the PBN could potentially have a role in alcohol consumption.

### NTS^CeA^ projection to the parabrachial nucleus (PBN) is reinforcing

Prior to investigating the role of the *NTS*^CeA➔PBN^ on consummatory behavior, we assayed the behavioral effects of pathway stimulation on measures of anxiety-like behavior and appetitive/aversive behavior. Consistent with the lack of effect on anxiety-like behavior noted with *NTS*^CeA^::casp mice, 20 Hz optical activation of the *NTS*^CeA➔PBN^::ChR2 pathway did not alter time spent in the center of an open field (Fig 8A; Unpaired t-test: t(7)=1.163, p=0.2830). Stimulation of the *NTS*^CeA➔PBN^ projection also failed to impact behavior in the elevated plus maze either in open arm entries (Fig 8B; Two-way ANOVA: interaction F_(2,27)_=0.01082, p=0.9892; stimulation, F_(2,27)_=0.1085, p=0.8976; virus type, F_(1,27)_= 0.4477, p=0.5091) or in time spent in the open arm (Fig C; interaction F_(2,27)_= 0.6265, p=0.5421; stimulation, F_(2,27)_= 3.034, p=0.0648; virus type, F_(1,27)_= 0.6867, p=0.4146), indicating that activating this pathway in naïve mice does not alter anxiety-like behaviors.

**Fig 8:**
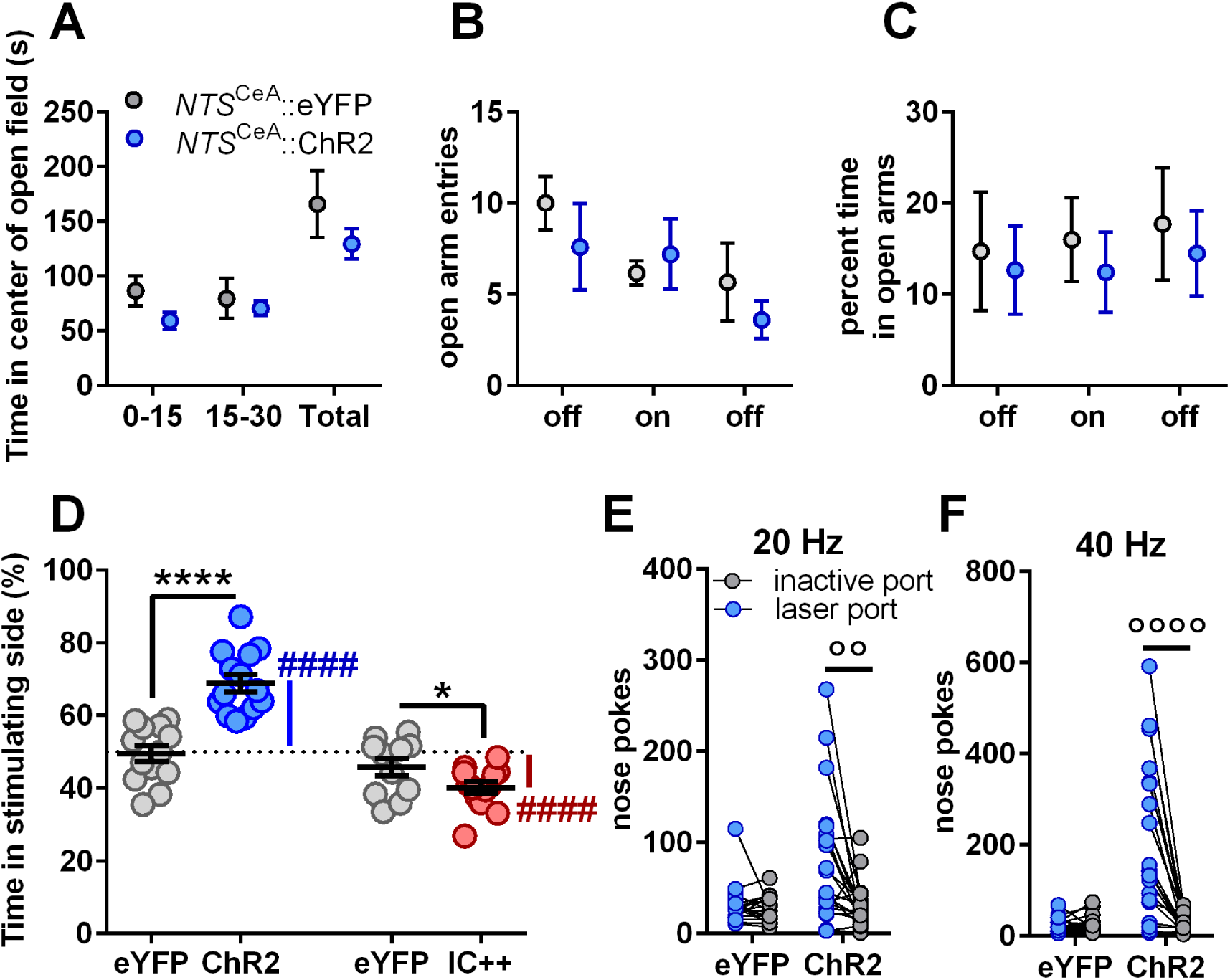
NTS^CeA->PBN^ optogenetic stimulation confers positive valence. (a) Optical stimulation in *NTS*^CeA➔PBN^::ChR2 (n=5) and *NTS*^CeA➔PBN^::eYFP mice (n=4) did not change time spent in the center of an open field. (**b**) Optical stimulation in *NTS*^CeA➔PBN^::ChR2 (n=5) and *NTS*^CeA➔PBN^::eYFP mice (n=6) did not impact either entries into or (**c**) time spent in the open arms of the elevated-plus maze. (**d**) *NTS*^CeA➔PBN^::ChR2 mice (n=14) spent significantly more time in the stimulation (20 Hz) side in a real-time place preference assay than *NTS*^CeA➔PBN^::eYFP mice (n=13), whereas *NTS*^CeA➔PBN^::IC++ mice (n=13) spent significantly less time in the stimulation side of this assay than *NTS*^CeA➔PBN^::eYFP controls (n=11). (**e**) *NTS*^CeA➔PBN^::ChR2 mice (n=18) nosepoked for 5 seconds of laser stimulation at both 20Hz and (**f**) 40 Hz stimulation whereas *NTS*^CeA➔PBN^::eYFP mice (n=22) did not. Unpaired t-test: *p≤0.05, **p<0.01, ****p<0.0001, One-sample t-test difference from 50%: ^####^p<0.0001, Bonferroni-corrected paired t-test: °°p<0.001, °°°°p<0.0001.

To probe if stimulation of the *NTS*^CeA➔PBN^ pathway altered affective valence, we examined response to photostimulation in the real-time place preference (RTPP) assay. Photo-stimulation of these fibers at 20 Hz induced a significant RTPP in *NTS*^CeA➔PBN^::ChR2-eYFP mice, but not in *NTS*^CeA➔PBN^::eYFP controls (Fig 8D; Unpaired t-test: t(25)=6.128, p<0.0001) suggesting that these neurons convey positive valence. We also wanted to confirm whether time spent in the stimulation side was significantly different from chance and found that this was the case for *NTS*^CeA➔PBN^::ChR2-eYFP mice (One-sample t-test: control: t(12)=0.2835, p=0.7817, ChR2-eYFP: t(13)=8.183, p<0.0001). To inhibit the terminals of *NTS*^CeA^ neurons in the PBN we expressed the blue light activated chloride channel IC++ (Berndt et al., 2016). We validated that viral IC++ expression in *NTS*^CeA^ neurons prevented action potential firing *ex vivo* (data not shown). When we expressed IC++ in the CeA and placed fibers in the PBN (*NTS*^CeA➔PBN^::IC++-eYFP), mice showed a mild aversion to inhibition of the projection (constant light stimulation, Fig 8D; Unpaired t-test: t(22)=2.071, p=0.0503). Congruently, we found that the *NTS*^CeA➔PBN^::IC++-eYFP animals but not the *NTS*^CeA➔PBN^::eYFP controls behaved significantly differently from chance (One-sample t-test: control: t(10)=1.774, p=0.1064, IC++-eYFP: t(12)=6.180, p<0.0001). Finally, *NTS*^CeA➔PBN^::ChR2 mice performed optical intracranial self-stimulation (oICSS) for 20 Hz (Fig 8E; Bonferroni corrected t-test active vs active port: control t(34)=0.930211, p=0.35882; ChR2 t(42)=3.19163, p=0.00268) as well as 40 Hz stimulation (Fig 8F; Bonferroni corrected t-test active vs active port: control t(34)=0.0708983, p=0.943894; ChR2 t(42)=4.61353, p =0.00004), demonstrating that activation of this pathway is intrinsically reinforcing. These data suggest that the *NTS*^CeA➔PBN^ pathway may bidirectionally modulate reward seeking behavior.

### Stimulation of the NTS^CeA➔PBN^ projection promotes consumption of palatable fluids

We next examined the impact of photostimulation on the consumption of a variety of fluids in *NTS*^CeA➔PBN^::ChR2 mice. As schematized in Figure 9A, mice were habituated to the chamber for 4 days and allowed to consume the test fluid for 3 hours each day. Over the subsequent 4 days mice received 2 days of optical stimulation (non-contingent on the mouse’s location) in 5 min cycles alternated with 2 days without stimulation, again for 3 hours each day. Importantly, mice had food and water *ad lib* during the entire course of the experiment, thus were not especially motivated to eat or drink.

**Fig 9:**
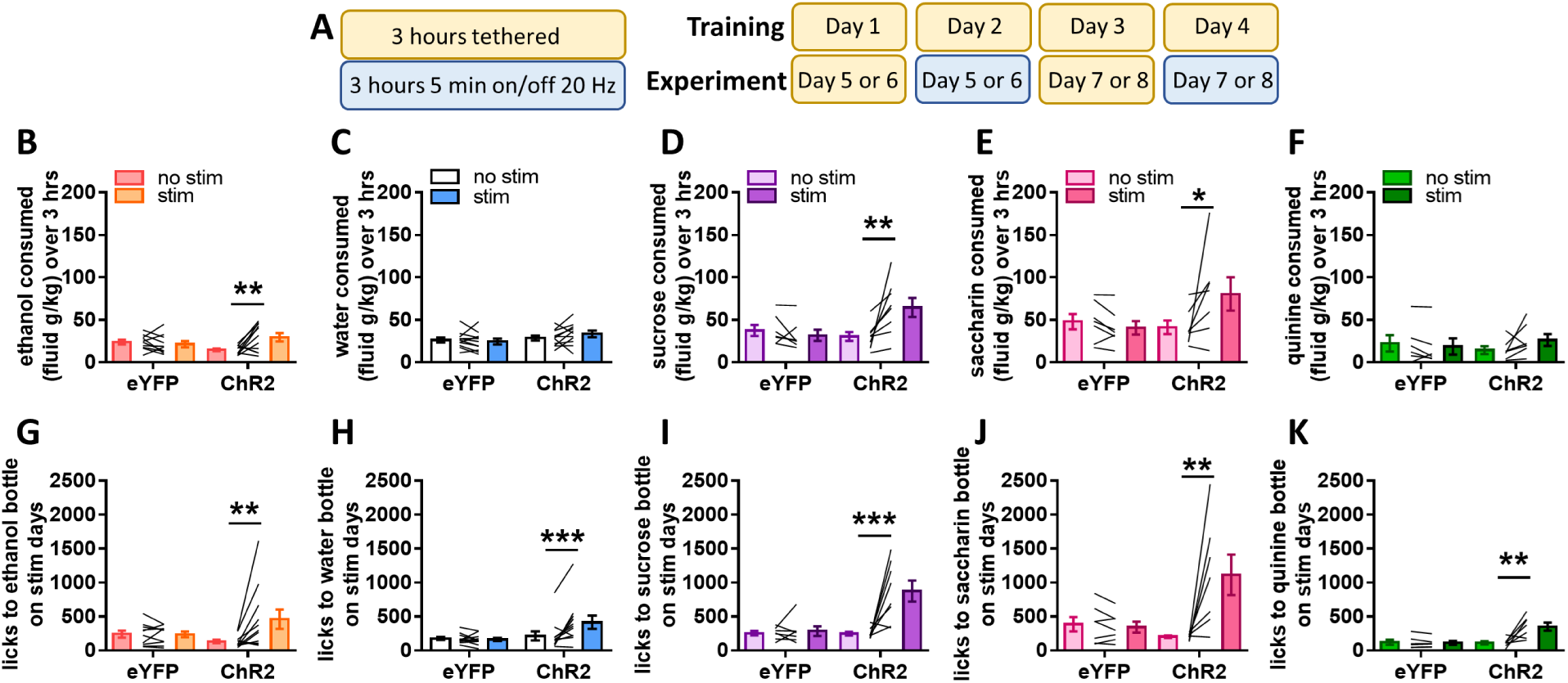
NTS^CeA->PBN^ optogenetic stimulation promotes consumption of rewarding fluids. (**a**) Schematic of optogenetic drinking paradigm. (**b**) *NTS*^CeA➔PBN^::ChR2 mice (n=11) drank significantly more ethanol (6% w/v) on stimulation days, while *NTS*^CeA➔PBN^::eYFP mice (n=10) were unaffected by stimulation. (**c**) *NTS*^CeA➔PBN^::ChR2 (n=12) and *NTS*^CeA➔PBN^::eYFP mice (n=11) drank similar amounts of water and this consumption was unaffected by optical stimulation. (**d**) *NTS*^CeA➔PBN^::ChR2 (n=7) mice drank significantly more sucrose (1% w/v) on stimulation days, while *NTS*^CeA➔PBN^::eYFP mice (n=7) were unaffected by optical stimulation. (**e**) *NTS*^CeA➔PBN^::ChR2 (n=7) mice drank significantly more saccharin (0.003% w/v) on stimulation days, while *NTS*^CeA➔PBN^::eYFP mice (n=7) were unaffected by optical stimulation. (**f**) *NTS*^CeA➔PBN^::ChR2 (n=7) and *NTS*^CeA➔PBN^::eYFP mice (n=6) drank similar amounts of quinine (100 μM), and this consumption was unaffected by optical stimulation. (**g-k**) *NTS*^CeA➔PBN^::ChR2 mice licked the bottle significantly more during stimulation epochs than during non-stimulation epochs in all conditions. Bonferroni-corrected paired t-test: *p<0.05, **p<0.01, ***p<0.001.

*NTS*^CeA➔PBN^::ChR2 and *NTS*^CeA➔PBN^::eYFP mice showed similar levels of ethanol drinking during habituation days (data not shown). We found that optical stimulation of the *NTS*^CeA➔PBN^ pathway increased consumption of 6% ethanol (Fig 9B; Two-way ANOVA: interaction F_(1,19)_=7.363, p=0.0138; virus type, F_(1,19)_=0.01524, p=0.9031; stimulation, F_(1,19)_=3.665, p=0.0707; Bonferroni-corrected t-test: control t(19)=0.5520, p>0.9999; ChR2 t(19)=3.353, p=0.0067) as compared to non-stimulation days, whereas stimulation of *NTS*^CeA➔PBN^::eYFP mice did not alter ethanol consumption. Examining only the days that the mice received stimulation, *NTS*^CeA➔PBN^::ChR2 mice licked the bottle significantly more during the 5-min laser on versus laser off phases (Fig 9G; Two-way ANOVA: interaction F_(1,19)_=6.117, p=0.0230; virus type, F_(1,19)_=0.3760, p=0.5470; stimulation, F_(1,19)_=5.890, p=0.0253; Bonferroni-corrected t-test: control t(19)=0.03198, p>0.9999; ChR2 t(19)=3.3551, p=0.0043).

We next sought to determine whether this increase in ethanol consumption was due to a generalized increase in liquid consumption, or an ethanol-specific phenotype. In mice given *ad libitum* food and water, we performed the same experimental paradigm as above, but with water instead of ethanol. Stimulation of *NTS*^CeA➔PBN^::ChR2 mice did not significantly alter water consumption (Fig 9C; Two-way ANOVA: interaction F_(1,21)_=1.901, p=0.1825; virus type, F_(1,21)_=0.5904, p=0.4508; stimulation, F_(1,21)_=0.2757, p=0.6051). Interestingly, however, on the stimulation days, the *NTS*^CeA➔PBN^::ChR2 mice engaged the water bottle more during the 5 minute laser stim epochs than the 5 minute non-stim epochs (Two-way ANOVA: interaction F_(1,21)_=8.591, p=0.0080; virus type, F_(1,21)_=2.397, p=0.1365; stimulation, F_(1,21)_=6.215, p=0.0211; Bonferroni-corrected t-test: control t(21)=0.3033, p>0.9999; ChR2 t(21)=3.922, p=0.0016). These results suggest that our optogenetic experiments are not manipulating a general fluid consumption pathway, like the neighboring *NTS*^LH^ neuron population (Kurt et al., 2018), but perhaps a more selective circuit for which the appetitive properties of the available fluid is important.

To determine whether stimulation of the *NTS*^CeA➔PBN^ projection would increase consumption of other palatable fluids, we performed the same experimental paradigm in the presence of 1% sucrose or 0.03% saccharin. *NTS*^CeA➔PBN^::ChR2 mice consumed significantly more sucrose solution on stimulation days (Fig 9D; Two-way ANOVA: interaction F_(1,12)_=10.23, p=0.0077; virus type, F_(1,12)_=2.584, p=0.1340; stimulation, F_(1,12)_=5.597, p=0.0357; Bonferroni-corrected t-test: control t(12)=0.5884, p>0.9999; ChR2 t(12)=3.934, p=0.0040), and licked the bottle significantly more during stimulation epochs (Fig 9I; Two-way ANOVA: interaction F_(1,12)_=15.92, p=0.0018; virus type, F_(1,12)_=13.89, p=0.0029; stimulation, F_(1,12)_=18.65, p=0.0010; Bonferroni-corrected t-test: control t(12)=0.2322, p>0.9999; ChR2 t(12)=5.875, p=0.0002). *NTS*^CeA➔PBN^::ChR2 mice also consumed significantly more saccharin solution on stimulation days (Fig 9E;Two-way ANOVA: interaction F_(1,12)_=4.946, p=0.0461; virus type, F_(1,12)_=1.490, p=0.2457; stimulation, F_(1,12)_=2.312, p=0.1543; Bonferroni-corrected t-test: control t(12)=0.4975, p>0.9999; ChR2 t(12)=2.648, p=0.0425), and licked the bottle more during stimulation epochs (Fig 9J;Two-way ANOVA: interaction F_(1,12)_=9.380, p=0.0099; virus type, F_(1,12)_=2.974, p=0.1103; stimulation, F_(1,12)_=7.776, p=0.0164; Bonferroni-corrected t-test: control t(12)=0.1938, p>0.9999; ChR2 t(12)=4.137, p=0.0028), indicating that the increase in consumption is not dependent on the caloric content of the solution.

We then performed the same experiment using a 100 μM quinine solution to determine whether *NTS*^CeA➔PBN^ stimulation would affect consumption of negative valence tastants. Stimulation failed to increase quinine drinking on stim vs no stim days (Fig 9F; Two-way ANOVA: interaction F_(1,11)_=3.137, p=0.1042; virus type, F_(1,11)_=0.0003, p=0.9859; stimulation, F_(1,11)_=0.8933, p=0.3649), but increased licking during stim vs no stim epochs (Fig 9K; Two-way ANOVA: interaction F_(1,11)_=9.798, p=0.0096; virus type, F_(1,11)_=7.165., p=0.0215; stimulation, F_(1,11)_=8.360., p=0.0147; Bonferroni-corrected t-test: control t(11)=0.1628, p>0.9999; ChR2 t(11)=4.432, p=0.0020). Taken together, these data suggest that stimulation of the NTS-CeA to PBN pathway increases consumption of rewarding fluids.

We next re-analyzed the videos from 3 of the consumption experiments (water-neutral, sucrose-palatable, and quinine-aversive) in order to validate the automated licking results. This was particularly important due to the discrepancy between the findings that *NTS*^CeA➔PBN^ stimulation increases bottle interaction regardless of fluid content (Fig 9G-K), but only increases consumption on days when the bottle contains a palatable/rewarding fluid (Fig 9B-F). We hand scored bottle-licking behavior and found that indeed *NTS*^CeA➔PBN^::ChR2 animals licked the bottle more on average during laser stimulation-on epochs regardless of whether the bottle contained water (Fig 10A; Two-way ANOVA: interaction F_(1,19)_=10.14, p=0.0049; virus type, F_(1,19)_=6.001, p=0.0242; stimulation, F_(1,19)_=10.52, p=0.0043; Bonferroni-corrected t-test: control t(19)=0.04096, p>0.9999; ChR2 t(19)=4.658, p=0.0003), sucrose (Fig 10B; Two-way ANOVA: interaction F_(1,13)_=10.27, p=0.0069; virus type, F_(1,13)_=11.80, p=0.0044; stimulation, F_(1,13)_=11.80, p=0.5824; Bonferroni-corrected t-test: control t(13)=0.1570, p>0.9999; ChR2 t(13)=4.860, p=0.0006) or quinine (Fig 10C; Two-way ANOVA: interaction F_(1,11)_=0.6329, p=0.0287; virus type, F_(1,11)_=0.2777, p=0.6087; stimulation, F_(1,11)_=4.107, p=0.0676; Bonferroni-corrected t-test: control t(11)=0.3333, p>0.9999; ChR2 t(11)=3.343, p=0.0131). These data reinforce the idea that stimulation of the *NTS*^CeA➔PBN^ pathway increases licking behavior, but that the relationship between licking behavior and fluid consumption is not 1:1.

**Fig 10:**
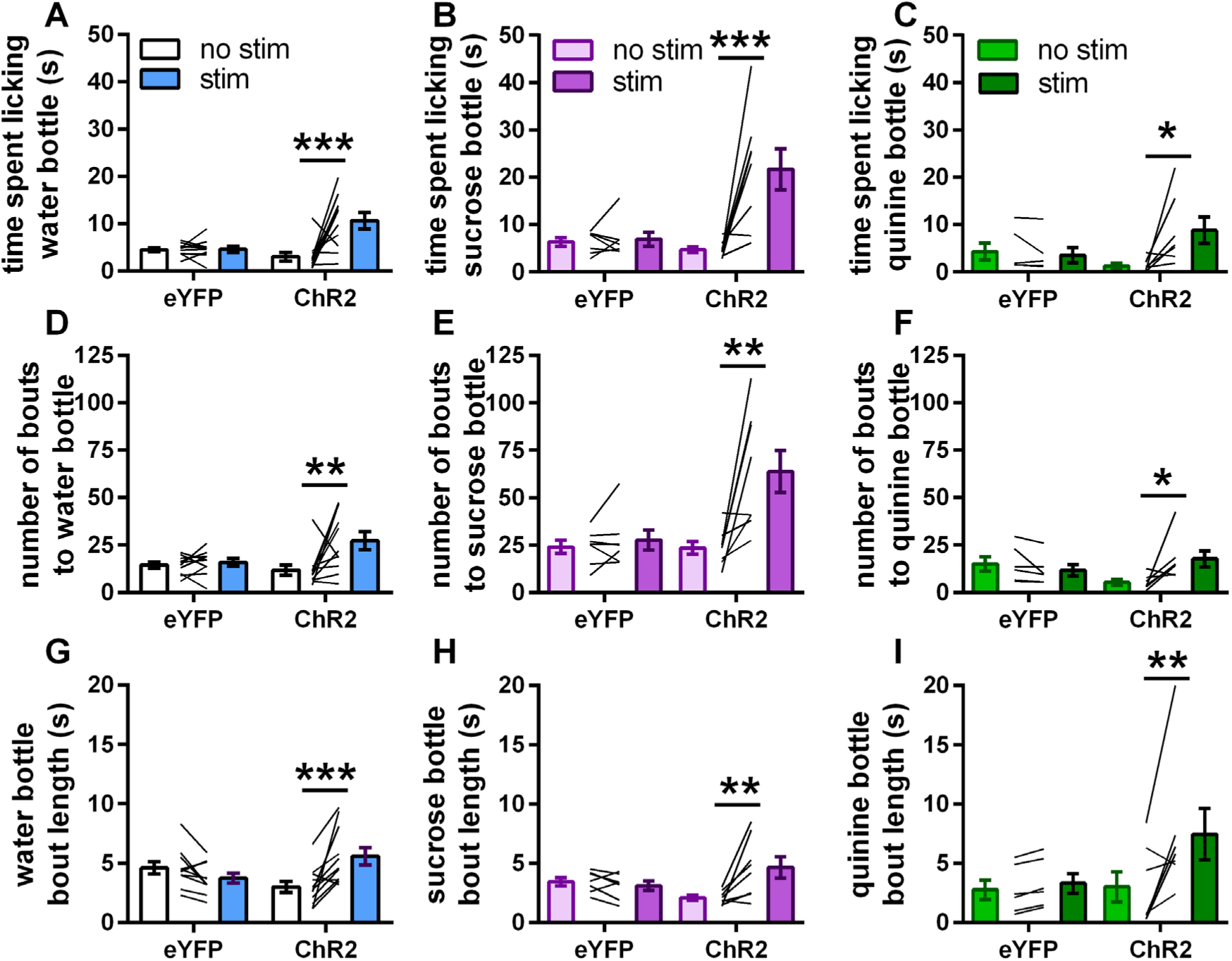
NTS^CeA->PBN^ optogenetic stimulation increases licking by increasing both bout length and number. (**a-c**) *NTS*^CeA➔PBN^::ChR2 mice spent more time licking the bottle during laser stimulation regardless of whether the bottle contained (**a**) water, (**b**) sucrose, or (**c**) quinine. Value is the average time spent licking across laser on-off epochs. (**d-f**) *NTS*^CeA➔PBN^::ChR2 mice had a higher number of drinking bouts regardless of whether the bottle contained (**d**) water, (**e**) sucrose, or (**f**) quinine. (**g-i**) Laser stimulation increased average bout length in *NTS*^CeA➔PBN^::ChR2 mice regardless of whether the bottle contained (g) water, (**h**) sucrose, or (**i**) quinine. Bonferroni-corrected paired t-test: *p<0.05, **p<0.01, ***0.001.

Previous work exploring the *Htr2a*^CeA➔PBN^ projection in consumption showed that optogenetic stimulation of this pathway increased the duration of feeding bouts (Douglass et al., 2017). We thus examined whether the number and/or duration of drinking bouts were affected with stimulation of the *NTS*^CeA➔PBN^ pathway. When we examined the number of drinking bouts across the whole 3 hours, we found that *NTS*^CeA➔PBN^::ChR2 animals initiated significantly more bouts during laser-on epochs regardless of whether the bottle contained water (Fig 10D; Two-way ANOVA: interaction F_(1,19)_=4.643, p=0.0442; virus type, F_(1,19)_=2.062, p=0.1673; stimulation, F_(1,19)_=6.764, p=0.0176; Bonferroni-corrected t-test: control t(19)=0.3081, p>0.9999; ChR2 t(19)=3.446, p=0.0054), sucrose (Fig 10E;Two-way ANOVA: interaction F_(1,13)_=7.675, p=0.0159; virus type, F_(1,13)_=6.283, p=0.0263; stimulation, F_(1,13)_=10.95, p=0.0057; Bonferroni-corrected t-test: control t(13)=0.3687, p>0.9999; ChR2 t(13)=4.45, p=0.0013) or quinine (Fig 10F; Two-way ANOVA: interaction F_(1,11)_=7.126, p=0.0218; virus type, F_(1,11)_=0.2517, p=0.6258; stimulation, F_(1,11)_=2.273, p=0.1598; Bonferroni-corrected t-test: control t(11)=0.7916, p=0.8907; ChR2 t(11)=3.074, p=0.0212). We found that stimulation also increased average bout length in *NTS*^CeA➔PBN^::ChR2 mice in the water (Fig 10G; Two-way ANOVA: interaction F_(1,19)_=16.03, p=0.0008; virus type, F_(1,19)_=0.03605, p=0.8514; stimulation, F_(1,19)_=3.896, p=0.0631; Bonferroni-corrected t-test: control t(19)=1.403, p=0.3537; ChR2 t(19)=4.331, p=0.0007), sucrose (Fig 10H; Two-way ANOVA: interaction F_(1,13)_=9.659, p=0.0083; virus type, F_(1,13)_=0.02477., p=0.8774; stimulation, F_(1,13)_=5.637, p=0.0337; Bonferroni-corrected t-test: control t(13)=0.5022, p>0.9999; ChR2 t(13)=4.013, p=0.0030), and quinine conditions (Fig 10I; Two-way ANOVA: interaction F_(1,11)_=4.571, p=0.0558; virus type, F_(1,11)_=1.372, p=0.2663; stimulation, F_(1,11)_=7.532, p=0.0191; Bonferroni-corrected t-test: control t(11)=0.4132, p>0.9999; ChR2 t(11)=3.593, p=0.0084). Thus, our data demonstrate that even when total liquid consumption is not altered by stimulation (water/quinine), the stimulation of this pathway promotes multiple behaviors associated with the seeking of fluids.

*Stimulation of the NTS*^CeA➔PBN^ *projection fails to impact consumption of solid foods under most conditions* The PBN has a well-described role in appetite suppression (Carter et al., 2013). Indeed, recent work describing a CeA to PBN projection indicates that GABAergic input from the CeA can promote food consumption (Douglass et al., 2017). Suppression of PBN anorexigenic neuronal ensembles could explain the increase in palatable fluid consumption observed in the previous experiments. If this were the case, however, we would expect stimulation of the *NTS*^CeA➔PBN^ pathway to induce an overall increase in consumption, reflected in chow intake over this same period. Stimulation of the *NTS*^CeA➔PBN^ pathway failed to impact chow consumption in the presence of water (Fig 11A; Two-way ANOVA: interaction F_(1,21)_=0.03704, p=0.8492; virus type, F_(1,21)_=0.003276, p=0.9549; stimulation, F_(1,21)_=3.223, p=0.0870), sucrose (Fig 11B; Two-way ANOVA: interaction F_(1,12)_=1.981, p=0.1846; virus type, F_(1,12)_=0.8698, p=0.3694; stimulation, F_(1,12)_=0.1347, p=0.7200), saccharin (Fig 11C; Two-way ANOVA: interaction F_(1,12)_=0.008336, p=0.9288; virus type, F_(1,12)_=0.4687, p=0.5066; stimulation, F_(1,12)_=1.952, p=0.1876) or quinine (Fig 11D; Two-way ANOVA: interaction F_(1,11)_=0.02909, p=0.8677; virus type, F_(1,11)_=0.1673, p=0.6904; stimulation, F_(1,11)_=0.001504, p=0.9698). Surprisingly, in the presence of ethanol, however, *NTS*^CeA➔PBN^::ChR2 mice decreased chow consumption on days when they received stimulation (Fig 11E; Two-way ANOVA: interaction F_(1,22)_=4.313, p=0.0497; virus type, F_(1,22)_=0.5391, p=0.4705; stimulation, F_(1,22)_= 7.387, p=0.0126; Bonferroni-corrected t-test: control t(19)=0.1007, p>0.9999; ChR2 t(19)=2.956, p=0.0162). Taken as a whole these data indicate that the *NTS*^CeA➔PBN^ projection is involved with rewarding fluid intake as opposed to general consumption.

**Fig 11:**
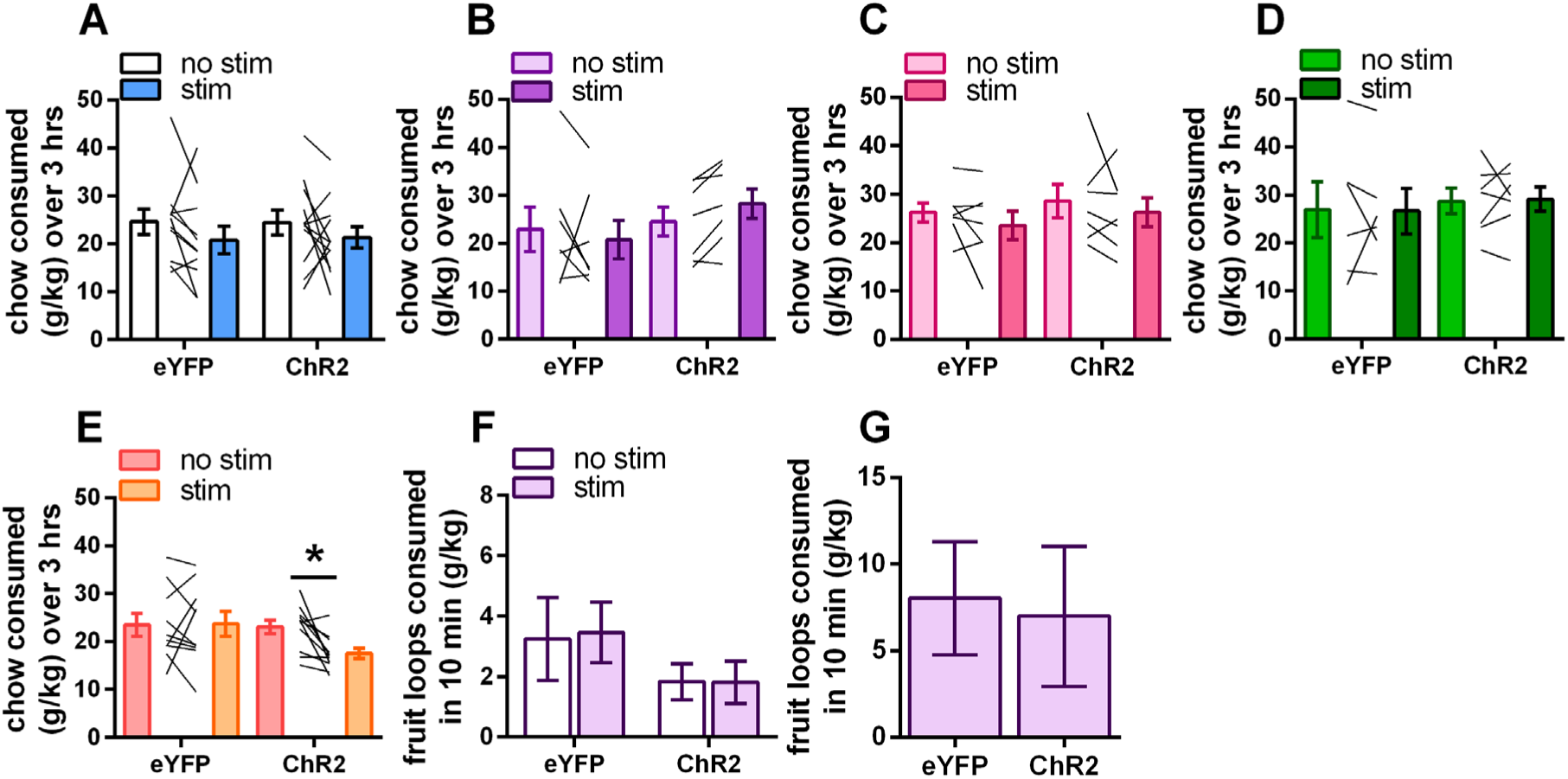
NTS^CeA->PBN^ optogenetic stimulation does not alter consumption of solid foods under most conditions. (**a-e**) Chow consumed during the optogenetic experiment outlined in Fig 9 in presence of (a) water, (**b**) sucrose, (**c**) saccharin, (**d**) quinine, and (**e**) ethanol. *NTS*^CeA➔PBN^::ChR2 and *NTS*^CeA➔PBN^::eYFP mice consumed similar amounts of chow during optogenetic stimulation. (**e**) *NTS*^CeA➔PBN^::ChR2 mice ate less chow on stimulation days when ethanol was present. (**f**) Stimulation failed to impact Froot Loop™ consumption during a 10-minute session regardless of whether the animals were sated (eYFP n=6, ChR2 n=7) or (**g**) following 24-hour food restriction (eYFP n=11, ChR2 n=14). Bonferroni-corrected paired t-test: *p<0.05.

Because optical stimulation of the *NTS*^CeA➔PBN^ promoted the consumption of sweet fluids, we then examined whether stimulation of this projection would impact consumption of a familiar sugary solid food. 2 days after homecage exposure to Froot Loops^TM^, *NTS*^CeA➔PBN^::ChR2 animals were allowed to consume Froot Loops^TM^ *ad lib* for 10 minutes. Optical stimulation of the *NTS*^CeA➔PBN^ did not impact Froot Loops^TM^ consumption (Fig 10F; Two-way ANOVA: interaction F_(1,11)_=0.01094, p=0.9186; virus type, F_(1,11)_=4.714, p=0.0527; stimulation, F_(1,11)_=0.007948, p=0.9306). In order to determine whether increasing the motivation to eat would perhaps reveal a role for this projection in palatable food consumption, we repeated this experiment following 24 hours of food restriction. Under these conditions stimulation failed to impact Froot Loops^TM^ consumption (Fig 10G; Unpaired t-test t(23)=0.7030, p=0.4891). Together, these data demonstrate a role for the *NTS*^CeA➔PBN^ projection in promoting the consumption of palatable fluids, disassociated from the CeA and PBN’s respective reported roles in solid food consumption.

## Discussion

The CeA regulates several behaviors associated with alcohol use disorders. The particular genetically defined cell types and circuits that mediate these behaviors, however, are poorly understood. Here we have shown that NTS-expressing neurons in the CeA contribute to voluntary ethanol consumption in non-alcohol dependent mice. Additionally, our data demonstrate that a subset of these neurons project to the PBN, that stimulation of this projection is positively reinforcing (supporting RTPP and oICSS), and leads to increased consumption of palatable fluids and ethanol.

### CeA neurotensin neurons in ethanol consumption

The CeA is well known to be engaged by ethanol consumption and is implicated in mediating both the negative and positive reinforcing properties of ethanol (Koob et al., 1998; Koob, 2015). In keeping with this, early studies found that of pharmacological inhibition of GABA_A_ receptors in (Hyytiä and Koob, 1995), and chemical lesions of (Möller et al., 1997), the CeA reduce ethanol consumption without affecting water consumption. Our data show that relatively low *in vivo* ethanol consumption can activate *Nts*^CeAL^ neurons (Fig 6F), and that selectively lesioning *NTS*^CeA^ neurons decreases ethanol intake and preference, without altering consumption of other fluids (Figs 3 and 5). Concordant with this finding, optogenetic stimulation of the *NTS*^CeA➔PBN^ projection increased ethanol consumption (Fig 9B), but again did not alter consumption of water or quinine solutions (Fig 9C,F). Future work will examine which aspects of *NTS*^CeA^ signaling, such as GABA, NTS, and/or other peptides, are responsible for these results.

Studies conducted in animals dependent on, or consuming binge quantities of, ethanol have identified CeA CRF signaling and CRF^CeA^ neurons as a locus of ethanol effects on GABA transmission (Nie et al., 2004; Lowery-Gionta et al., 2012; Pleil et al., 2015; Herman et al., 2016; de Guglielmo et al., 2019). In fact, a recent study from de Guglielmo *et al*. (2019) showed that inhibition of the *Crh*^CeA->BNST^ projection in ethanol-dependent rats decreased ethanol intake and symptoms of somatic withdrawal, illustrating the potential of these neurons to mediate negative reinforcing aspects of ethanol consumption. Our data and others (Kim et al., 2017; McCullough et al., 2018) indicate that *Nts*^CeA^ neurons are a subset of *Crh^CeA^*and *Crh1^CeA^* neurons, suggesting that other genetically-overlapping CeA projections may also be modulated by a history of ethanol consumption.

*Nts*^CeA^ neurons also have a partial overlap with *Pdyn*^CeA^ neurons. Dynorphin neurons in the CeA contribute to binge-drinking, a form of ethanol consumption that confers a high risk of developing alcohol use disorder (Anderson et al., 2019). We recently showed that dynorphin and NTS bi-directionally modulate synaptic inputs from the CeA to the BNST (Normandeau et al., 2018). This phenomenon may also be relevant to intra-CeA signaling, as well as CeA➔PBN projections, and provide yet another mechanism for ethanol-induced plasticity in this circuit. Because of these data, we hypothesize that multiple CeA populations, including the *NTS*^CeA➔PBN^ projection, may mediate early positive reinforcement and therefore could facilitate the transition into dependence. While we were surprised that manipulation of *NTS^CeA^* neurons did not alter anxiety-like behavior, we also hypothesize that these neurons may play different roles depending on the state of the animal (e.g. stress, dependence, intoxication, thirst).

### Ethanol consumption and appetite

We found that stimulation of the *NTS*^CeA->PBN^ pathway decreased food consumption when ethanol was available. Ethanol consumption and appetite have a complex relationship that has not been fully parsed (Cains et al., 2017), and food consumption may impact subjective perceptions of the effects of ethanol consumption (Caton et al., 2007). Previous *ex vivo* studies have shown that the CeA is a site of action for the pharmacological effects of both ghrelin and ethanol (Cruz et al., 2013), suggesting that this may be a site of interplay between appetite and ethanol. Due to limitations of our experimental design, we were not able to explore this finding, but believe that further work examining this relationship in the context of the *NTS*^CeA->PBN^ circuit is promising.

### CeA neurotensin neurons promote positive valence behaviors

There is a general hypothesis that the CeA has a role in amplifying motivation for reward-seeking but does not have a direct role in reward in and of itself. This is largely because nonspecific optical CeA stimulation increases responding for a laser-paired positive reinforcer and can shift preference towards a non-preferred paired outcome (Robinson et al., 2014; Warlow et al., 2017). However, this manipulation does not support intracranial self-stimulation behavior for unpaired stimulation. On the other hand, our results demonstrating that optical stimulation of the *NTS*^CeA➔PBN^ pathway is reinforcing is consistent with recent data showing that NTS+ neurons in the CeA promote positive valence (Kim et al., 2017). While Kim *et al*. divided the *NTS*^CeA^ population into two groups, mice performed nose-poking behavior for cell-body stimulation for both of these subpopulations.

Because the CeA is composed of a heterogenous population of neurons expressing multiple neuropeptides/signaling molecules, projecting both within the nucleus and across the brain, we suggest that stimulation of the CeA as a whole may obscure the role of specific projections or genetically-defined subtypes, particularly if they have reciprocal inhibitory connections within the CeA. In addition to Kim *et al*, other work in CeA➔PBN projections from genetically-defined subtypes, such as *Htr2a* (serotonin 2a receptor) and *Pnoc* (prepronociceptin), have shown that stimulation can support nose-poking behavior (Douglass et al., 2017; Hardaway et al., 2019). Another explanation may be that most of the experiments examining genetically-defined CeA populations have been conducted in mice, whereas studies stimulating the CeA as a whole have largely been performed in rats (however see de Guglielmo *et al*., 2019).

Our finding that stimulation of the *NTS*^CeA➔PBN^ projection can both promote positive valence behaviors and increase consummatory behaviors are at first counterintuitive. Indeed, much work elucidating the neural circuits of feeding has described circuits that promote consumption through negative valence signals encoding hunger and thirst states (Betley et al., 2015). However, we are not alone in describing an amygdala-to-PBN circuit fulfilling both of these criteria. Recent experiments describe a CeA *Htr2a*-containing population that promotes food consumption (Douglass et al., 2017), which may overlap with the *Nts* population (Kim et al., 2017; Torruella-Suarez data not shown). These circuits may underlie hedonic consumption, a form of consumption that has particular implications for the obesity epidemic (Lowe and Butryn, 2007).

### Palatable fluid consumption: implications for sweetened beverages

While we show here that ablation of *NTS*^CeA^ neurons failed to impact preference for sweet or bitter fluids, stimulation of the *NTS*^CeA➔PBN^ projection increased consumption of a variety of palatable fluids, and revealed a role for this neuronal population in palatable fluid consumption. Our results, however, are markedly different to other fluid circuits that have been described within relevant NTS-neuron and PBN circuity. *Oxtr*^PBN^ neurons appear to signal overall fluid satiation (Ryan et al., 2017), whereas stimulation of *NTS*^LH^ neurons increases fluid consumption, regardless of the identity of the available fluid (Kurt et al., 2018). In contrast, our data demonstrates that ablation of the *NTS*^CeA^ neurons does not alter gross fluid consumption. While we do not know the precise identity of the neurons in the PBN that receive input from the *NTS*^CeA^ neurons, future work to classify which population is inhibited by the *NTS*^CeA^ will undoubtedly be very informative as to how this circuit regulates the consumption of palatable fluids.

While the current obesity epidemic clearly has a variety of causes, sweetened beverages have emerged as an important target for both study and policy intervention by concerned government entities (Fowler et al., 2008; Malik et al., 2013; CDC, 2017). Interestingly, ethanol has a sweet taste component in both humans and C57BL/6J mice (Scinska et al., 2000; Blizard, 2007), which may account for why stimulation of the *NTS*^CeA➔PBN^ pathway promoted its consumption. In contrast, caspase ablation of the *NTS*^CeA^ neurons impaired ethanol consumption without affecting sucrose or saccharin preference, which, in conjunction with our results showing that sucrose consumption elevated *Fos* in the CeA_M_, suggests that there may be redundant circuitries that compensate for the drive to consume sweet beverages. Regardless, it is worth noting that consumption of alcoholic beverages by people almost always includes sweeteners. The connection between ethanol and sweet liquid consumption in our data presents an additional convergence between these consummatory behaviors, and future experiments will focus on understanding how sweet beverages and ethanol contribute to adaptations within this pathway.

Here we describe a genetically defined population of CeA neurons, *NTS*^CeA^, that are activated by ethanol drinking *in vivo,* and whose ablation impairs ethanol consumption and preference. Optical stimulation of the *NTS*^CeA➔PBN^ projection conferred a positive valence and increased consumption of rewarding fluids such as sweet flavored and ethanol solutions. Stimulation of this projection did not increase consumption of neutral or aversive fluids, impact consumption of solid food (with the intriguing exception of ethanol/chow choice) or affect anxiety-like behaviors. This work highlights the *NTS*^CeA➔PBN^ pathway as a fundamental circuit in promoting drinking behavior, and suggests that further examination of this pathway is relevant for the study of motivation to consume in the context of obesity and alcohol use disorders.

